# A CTCF-LDHB axis converts nuclear lactate availability into an H3K18 acylation switch that directs alternative splicing in hypoxic breast cancer

**DOI:** 10.64898/2026.10.27.762200

**Authors:** Parik Kakani, Abhisek Brahma, B Jithin, Rose Maria Peter, Abhisek Panda, Anamika Bansal, Om Prakash Kumar, Kaustubh Niyati, Abin Sahu, Atul Samaiya, Sanjeev Shukla

## Abstract

Lactate accumulating in hypoxic tumours lactylates histones, coupling metabolic state to chromatin. Why it differs between tumours of comparable glycolytic rate has remained unresolved. Here we show that nuclear, rather than total intracellular, lactate availability determines histone H3K18 lactylation in breast cancer, and that CCCTC-binding factor (CTCF) and lactate dehydrogenase B (*LDHB*) jointly set it. CTCF sustains glycolytic gene expression under hypoxia and drives *LDHB* in triple-negative breast cancer (TNBC), whereas a CTCF-ERα complex represses *LDHB* in non-TNBC. LDHB consumes lactate favouring H3K18 acetylation; where *LDHB* is low, CTCF-dependent MCT1 delivers lactate into the nucleus and sustains lactylation. At a CTCF site downstream of *TACC3* exon 3, an invariant polymerase II pause couples kinetic and adaptor control of splicing, with the mark deciding the outcome: H3K18la recruits SMARCA4 and hnRNPA1 to skip the exon, generating an isoform that drives EMT, whereas H3K18ac recruits TRIM33 and SRSF10 to include it, generating one that confers stemness. Inhibiting LDHB restores exon skipping and suppresses tumour growth in vivo. Nuclear lactate availability therefore acts as a metabolic switch that converts hypoxic stress into subtype-specific splicing decisions and identifies LDHB as a tractable target in TNBC.

## Introduction

Breast cancer remains a leading cause of morbidity and mortality worldwide despite advances in early detection, surgery, chemotherapy and targeted therapeutics ^1^. The major contributing factors are captured in the hallmarks of cancer, among which metabolism plays a prominent role, reflected in a now substantial body of work characterizing cancer as, in part, a systemic metabolic disease ^2^. To sustain a high proliferation rate under hypoxia, cancer cells undergo extensive metabolic reprogramming, shifting from oxidative phosphorylation towards accelerated aerobic glycolysis (the Warburg effect), characterized by elevated glucose uptake, high glycolytic flux and excessive lactate secretion ^3^. Although this pathway yields less ATP than oxidative metabolism, it supplies rapidly proliferating cells with glycolytic intermediates for nucleotide, amino acid and lipid biosynthesis. These features are not uniform across breast cancer subtypes. Triple-negative breast cancer (TNBC), defined by the absence of estrogen (ER) and progesterone (PR) receptors and of HER2 amplification, exhibits a more pronounced glycolytic phenotype than ER-positive luminal tumors, with higher glucose uptake and lactate production. While glycolytic enzymes are active across breast cancer subtypes, TNBC displays significant upregulation of key regulators such as GLUT1, PFKFB3, and LDHA. This results in hyperactive lactate production and export, which acidifies the tumor microenvironment and thereby promotes invasion and angiogenesis ^4,5^. Recent work has established lactate as more than a glycolytic end product: it functions as a signalling molecule that links metabolic reprogramming to chromatin regulation ^6^ ^7^.

Hypoxia-inducible factor 1 alpha (HIF-1α) and lactate levels alone do not account for the extensive promoter accessibility observed under hypoxia. A substantial literature implicates chromatin modifiers in the resulting differential gene expression ^8^. CCCTC-binding factor (CTCF) is one such factor, and our previous work has shown that it performs multiple roles, including genome-wide redistribution and the consequent regulation of alternative splicing ^9–12^. We have previously shown that *CTCF* is upregulated under hypoxia and influences RNA polymerase II (Pol II) transcriptional dynamics ^9,12^. The chromatin adaptor model provides a framework for how local epigenomic features dictate splice site selection: CTCF acts as a functional speedbump, inducing a localized kinetic pause of Pol II elongation. This delay creates a window in which reader proteins recognize specific post-translational modifications near splice junctions, and these readers in turn recruit RNA-binding proteins such as splicing factors, thereby shaping the splicing outcome ^13^. How CTCF redistribution, the associated metabolic by-products and the resulting epigenetic modifications together regulate co-transcriptional alternative splicing in TNBC and non-TNBC under hypoxia remains largely unknown.

Here we show how CTCF-mediated promoter architecture and the contrasting metabolic output of the two subtypes together regulate co-transcriptional alternative splicing in TNBC and non-TNBC. We find that CTCF controls LDHB expression in TNBC, increasing the turnover of lactate to pyruvate and thence to acetyl-CoA, and that CTCF also regulates *SLC2A3* (GLUT3) and *PFKFB3*, driving lactate production under hypoxia. Bioenergetic assays (ECAR and OCR) revealed a marked contrast in lactate handling between the two subtypes, which we link to CTCF-dependent regulation of the lactate transporters *SLC16A1* (MCT1) and *SLC16A3* (MCT4). We then show how this disparate metabolic status regulates co-transcriptional alternative splicing of *TACC3* through CTCF together with H3K18la and H3K18ac and their associated readers and splicing factors, yielding a different predominant isoform in each subtype. The full-length *TACC3* isoform (TACC3 FL) supports cancer stemness and self-renewal in TNBC cells, whereas the TACC3 Δ3 isoform promotes the EMT programme in non-TNBC cells.

## Results

### CTCF drives hypoxia-induced glycolytic reprogramming, and CTCF depletion creates an energy-restricted state in breast cancer cells under hypoxia

To test whether hypoxia-induced glycolysis depends on CTCF, we examined the relationship between *CTCF* expression and glycolytic score in hypoxic cells using a published single-cell RNA-seq dataset (GSE165929).

We first extracted the epithelial malignant cells (**Figure S1A and S1B**) and stratified these malignant epithelial cells along a continuous hypoxia score derived from the Buffa gene-set signature. Across these cells, *CTCF* expression was positively and significantly associated with both the Buffa hypoxia score and the glycolytic score (**Figure 1A and S1C and S1D**). Given that hypoxia induces *CTCF* expression, as shown previously ^9,12^, these data prompted us to hypothesize that *CTCF* contributes to the increased glycolytic flux of hypoxic cells.

**Figure 1:**
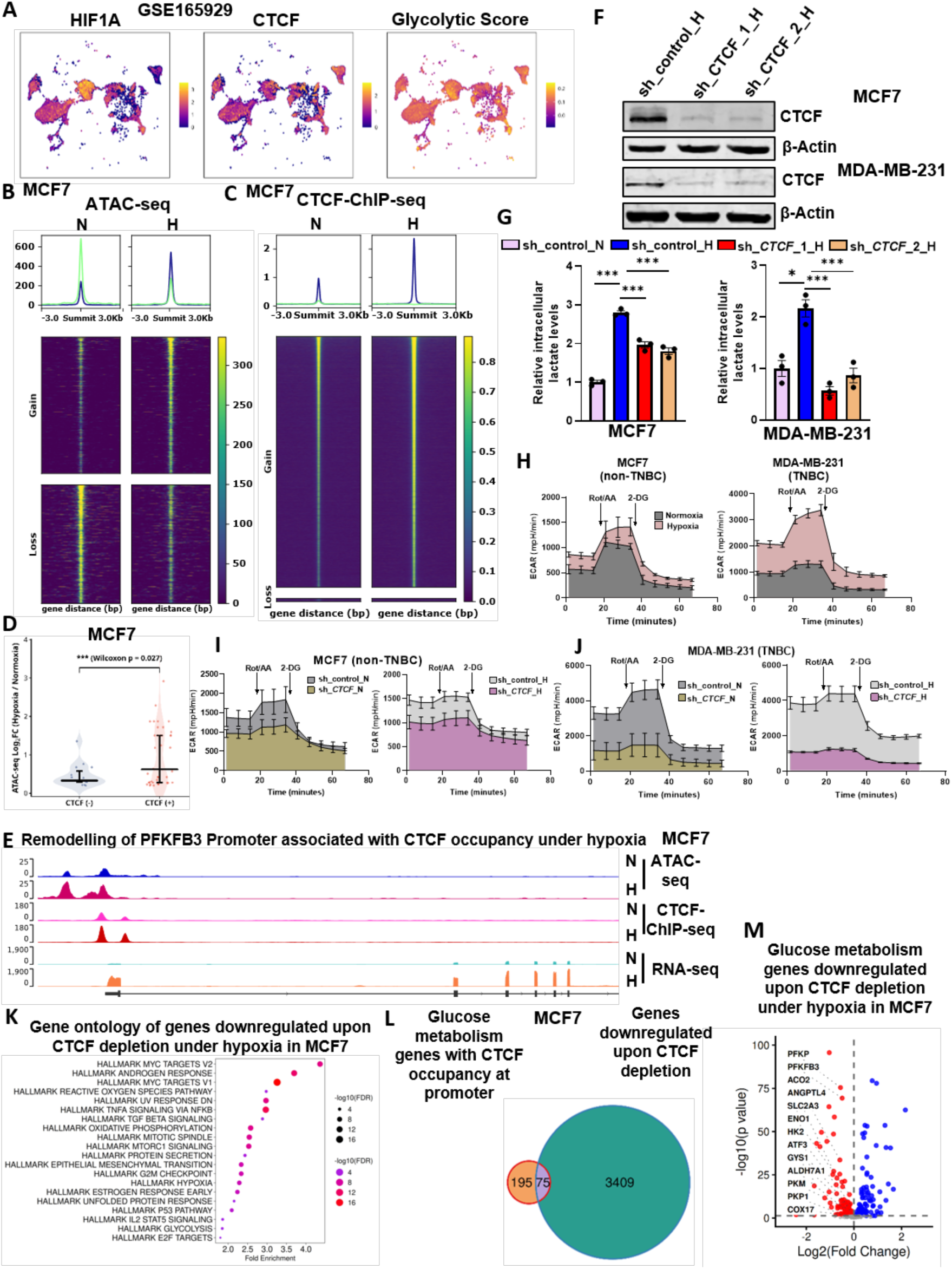
CTCF drives hypoxia-induced glycolytic reprogramming, and CTCF depletion creates an energy-restricted state in breast cancer cells under hypoxia. A) UMAP plot of breast epithelial cancerous cell populations stratified as continuous hypoxic state using Buffa score highlighting HIF1A, CTCF expression and glycolytic scores B) Differential binding analysis of ATAC signal between hypoxic and normoxic MCF7 cells inferred from ATAC-seq analysis C) Differential binding analysis of CTCF occupancy between hypoxic and normoxic MCF7 cells inferred D) Violin plot showing ATAC-seq gained accessibility (Hypoxia / Normoxia) at promoters of gene associated with glucose metabolism within a 2,000 bp window. CTCF(+) loci are associated with CTCF sites; CTCF(-) loci are not occupied by CTCF. Horizontal bars indicate median values E) PFKFB3 gene as an illustrative example of CTCF-associated chromatin opening under hypoxia, genome browser view of PFKFB3 loci of ATAC-seq, CTCF-ChIP-seq and RNA-seq in MCF7 cells under normoxia and hypoxia F) Immunoblot of CTCF in CTCF knockdown and shControl MCF7 cells under hypoxia G) Intracellular lactate level of sh_control and sh_CTCF MCF7 and MDA-MB-231 cells H) ECAR analysis of MCF7 and MDA-MB-231 cells under normoxia and hypoxia I-J) ECAR analysis of sh_control and sh_CTCF under hypoxia in I) MCF7 J) MDA-MB-231 cells. Cells were detected at basal level, followed by adding rotenone antimycin a (Rot/AA, 50μM) and 2-DG (50mM) K) Representative significantly enriched terms from the Gene Ontology of genes downregulated upon CTCF depletion under hypoxia in MCF7 cell line L) Venn diagrams showing the overlap between hypoxia-gained CTCF occupancy-related glucose metabolism gene and downregulated genes in CTCF depletion under hypoxia M) Volcano plot of differentially expressed glucose metabolism associated genes between sh_control and sh_CTCF under hypoxia and MCF7 cells. Red dots indicate downregulated genes with FDR <0.05. Data represent three technical replicates from pooled samples. Data are mean ± SEM, statistical significance was determined using Fischer’s exact test (A), one-way ANOVA with Tukey’s multiple comparisons test (F), one-sided Wilcoxon rank-sum test (D), *p < 0.05, **p < 0.01, and ***p < 0.001; n = 3 biological replicates.

To ask whether this association has a chromatin basis, we performed ATAC-seq in normoxic and hypoxic MCF7 cells (**Figure 1B**) and intersected the resulting accessibility profiles with CTCF ChIP-seq data in normoxic and hypoxic MCF7 cells (**Figure 1C, Table S1**) significant number of glucose metabolism genes that gained substantially more chromatin accessibility under hypoxia harbour CTCF at their promoters (**Figure 1D, Table S1**); *PFKFB3* is shown as a representative locus (**Figure 1E**).

Consistent with increased glycolytic flux, intracellular lactate levels rose under hypoxia in both cell lines, and *CTCF* depletion under hypoxia reduced intracellular lactate in each (**Figure 1F and G**). We next used Seahorse extracellular flux analysis to measure glycolytic capacity in TNBC (MDA-MB-231) and non-TNBC (MCF7) breast cancer cells under hypoxia (**Figure 1H**). Both lines showed increased extracellular acidification rate (ECAR) under hypoxia (**Figure 1H**), and both showed decreased ECAR upon *CTCF* depletion relative to control in normoxia as well as hypoxia (**Figure 1I and 1J**), indicating a substantial contribution of *CTCF* to hypoxia-mediated glucose metabolic programming. To define the genes through which CTCF acts under hypoxia, we depleted *CTCF* and performed RNA-seq (**Figure S1E**). Genes downregulated upon *CTCF* depletion under hypoxia were enriched for the glycolysis hallmark gene set (**Figure 1K**, **Table S1**). We then asked how many genes of the glucose metabolic network (glucose uptake, glycolysis and TCA cycle) carry CTCF at their promoters, and found that a substantial fraction do, with many of these downregulated upon *CTCF* depletion under hypoxia (**Figure 1L and 1M, Table S1**). Together, these data indicate that hypoxia-induced *CTCF* upregulation and redistribution of CTCF occupancy are required for induction of glucose uptake and glycolysis genes, and that *CTCF* depletion under hypoxia places breast cancer cells in an energy-restricted state.

### CTCF is required for optimal GLUT3 expression under hypoxia

To understand how *CTCF* supports hypoxia-induced glycolytic flux, we examined the RNA-seq data from *CTCF*-depleted hypoxic cells in more detail. The hypoxia-induced glucose uptake gene *GLUT3* (SLC2A3) showed reduced expression upon CTCF depletion under hypoxia (**Figure 2A, Table S1**). The glycolytic genes *HK2, PFKFB3* and *PFKP* were also reduced upon CTCF depletion in hypoxic breast cancer cells (**Figure 2A, Table S1**).

**Figure 2:**
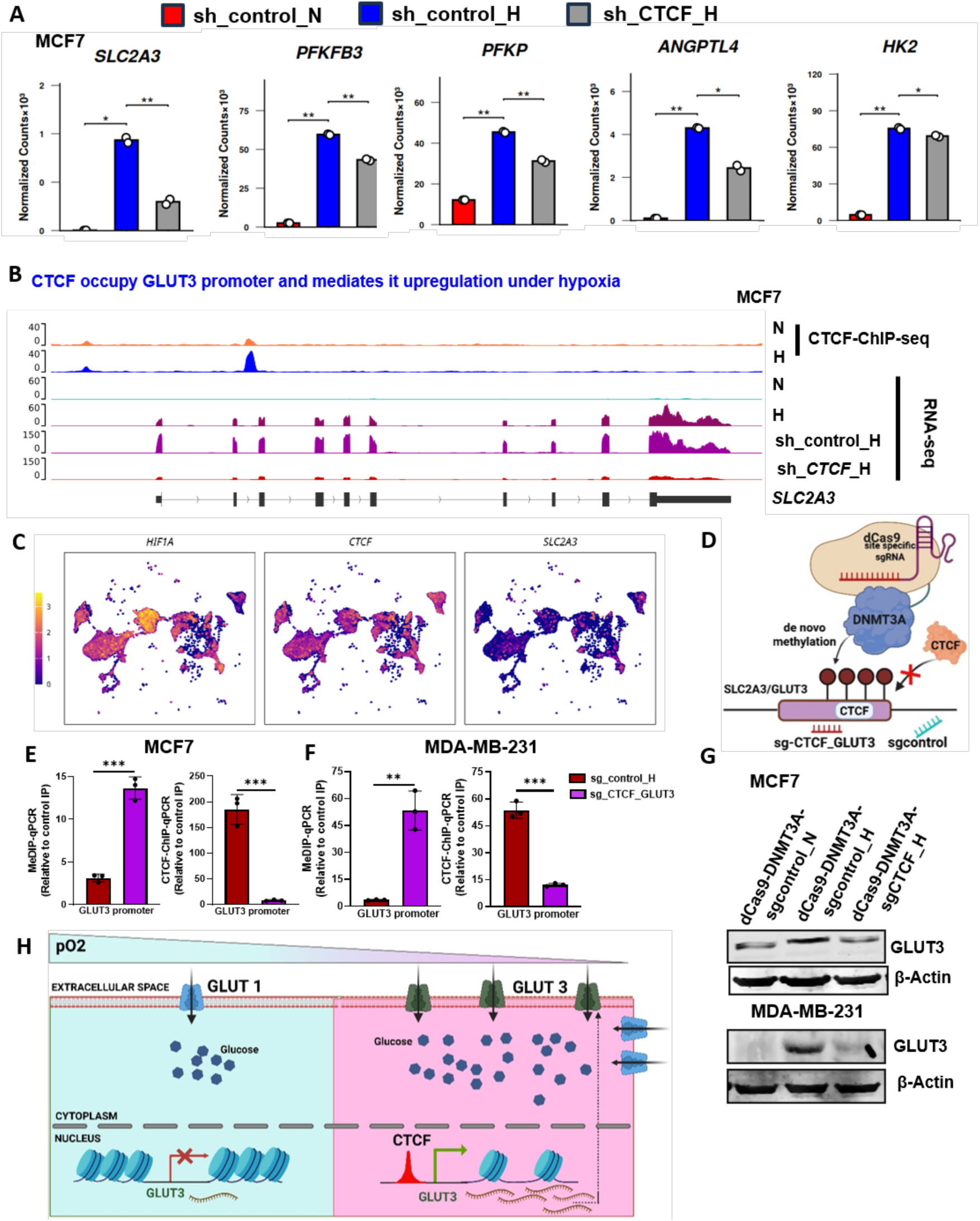
CTCF is required for optimal GLUT3 expression under hypoxia. A) mRNA expression of glycolysis genes in sh_control under normoxia and hypoxia and sh_CTCF under hypoxia in MCF7 cells inferred from CTCF depletion RNA-seq B) genome browser view of SLC2A3 loci of CTCF-ChIP-seq and RNA-seq under normoxia and hypoxia and of RNA-seq in shcontrol and shCTCF under hypoxia in MCF7 cells C) UMAP plot of breast epithelial cancerous cell populations stratified as continuous hypoxic state using Buffa score highlighting CTCF expression and SLC2A3 expression D) Schematic representation of targeting the CTCF motif present in the SLC2A3/GLUT3 promoter region by dCas9-DNMT3A with specific sgRNA against CTCF to maintain methylation and suppress CTCF occupancy E-G) E and F) MeDIP-qPCR and CTCF-ChIP-qPCR on GLUT3 promoter E) MCF7 and F) MDA-MB-231 (G) immunoblot of GLUT3 in MCF7 and MDA-MB-231cells transfected with dCas9-DNMT3A-sgCTCF in comparison to the sgcontrol under hypoxia (H) Schematic showing hypoxia-driven epigenetic and transcriptional regulation of GLUT3 expression in breast cancer. Data represent three technical replicates from pooled samples. Data are mean ± SEM, statistical significance was determined using unpaired, two-tailed Welch’s Two-Sample t-test (A), Co-expression was assessed using Fisher’s exact test (odds ratio [OR] = 1.80, p < 2.2 × 10⁻¹⁶) (C), unpaired two-tailed Student’s t-tests (F and G), *p < 0.05, **p < 0.01, and ***p < 0.001; n = 3 biological replicates.

To determine how CTCF regulates *GLUT3* under hypoxia, we analysed CTCF ChIP-seq and RNA-seqfrom normoxic and hypoxic cells with the CTCF-depleted RNA-seq data under hypoxia. CTCF occupies the *GLUT3* promoter, its expression increased under hypoxia and *GLUT3* expression falls upon CTCF depletion under hypoxia, consistent with direct regulation of *GLUT3* by CTCF under hypoxia (**Figure 2B, Table S1**).

To extend this observation to patient data, we examined *CTCF* and *GLUT3* in the single-cell RNA-seq dataset described above and found a highly significant positive correlation between *CTCF* and *GLUT3* expression that tracked with the hypoxic state of the cells (**Figure 2C and Figure S1F**).

To test whether CTCF binding is required, we used the dCas9-DNMT3A epigenome editing system, which deposits de novo 5-methylcytosine (5mC) and thereby displaces CTCF, as described previously ^9^ (**Figure 2D**). dCas9-DNMT3A-sgCTCF directed to the CTCF binding site at the *GLUT3* promoter increased 5mC and reduced CTCF occupancy in both cell lines relative to dCas9-DNMT3A-sgcontrol under hypoxia (**Figure 2E and F**). *GLUT3* expression was correspondingly reduced in dCas9-DNMT3A-sgCTCF-GLUT3 cells relative to dCas9-DNMT3A-sgcontrol under hypoxia (**Figure 2G**).

Together, these data show that hypoxia-induced *GLUT3* expression requires CTCF, and that CTCF sustains the energy state of hypoxic cells by supporting glucose uptake and the downstream glycolytic flux that generates lactate (**Figure 2H**).

### CTCF drives LDHB expression in TNBC, whereas a CTCF-ERα complex represses LDHB in non-TNBC

To examine how CTCF shapes the broader metabolic state, we used Seahorse analysis to quantify mitochondrial respiration. Under hypoxia the oxygen consumption rate (OCR) fell markedly in MCF7 cells, whereas in MDA-MB-231 cells OCR was largely maintained (**Figure 3A**). *CTCF* depletion did not significantly change OCR in MCF7 cells (**Figure 3B**), but markedly reduced OCR in MDA-MB-231 cells under hypoxia (**Figure 3C**), indicating that CTCF supports mitochondrial respiration specifically in TNBC cells. To identify the underlying mechanism, we examined CTCF ChIP-seq and RNA-seq in MCF7 and MDA-MB-231 cells and found that CTCF occupies the *LDHB* promoter in both lines (**Figure 3D, Table S2**), confirmed by CTCF-ChIP-qPCR (**Figure 3E**), even though *LDHB* is differentially expressed and is markedly higher in MDA-MB-231 cells (**Figure 3F**). The *LDHB* promoter carried the repressive H3K27me2 mark in MCF7 cells, whereas the active H3K27ac mark occupied the *LDHB* promoter in MDA-MB-231 cells (**Figure 3G**).

**Figure 3:**
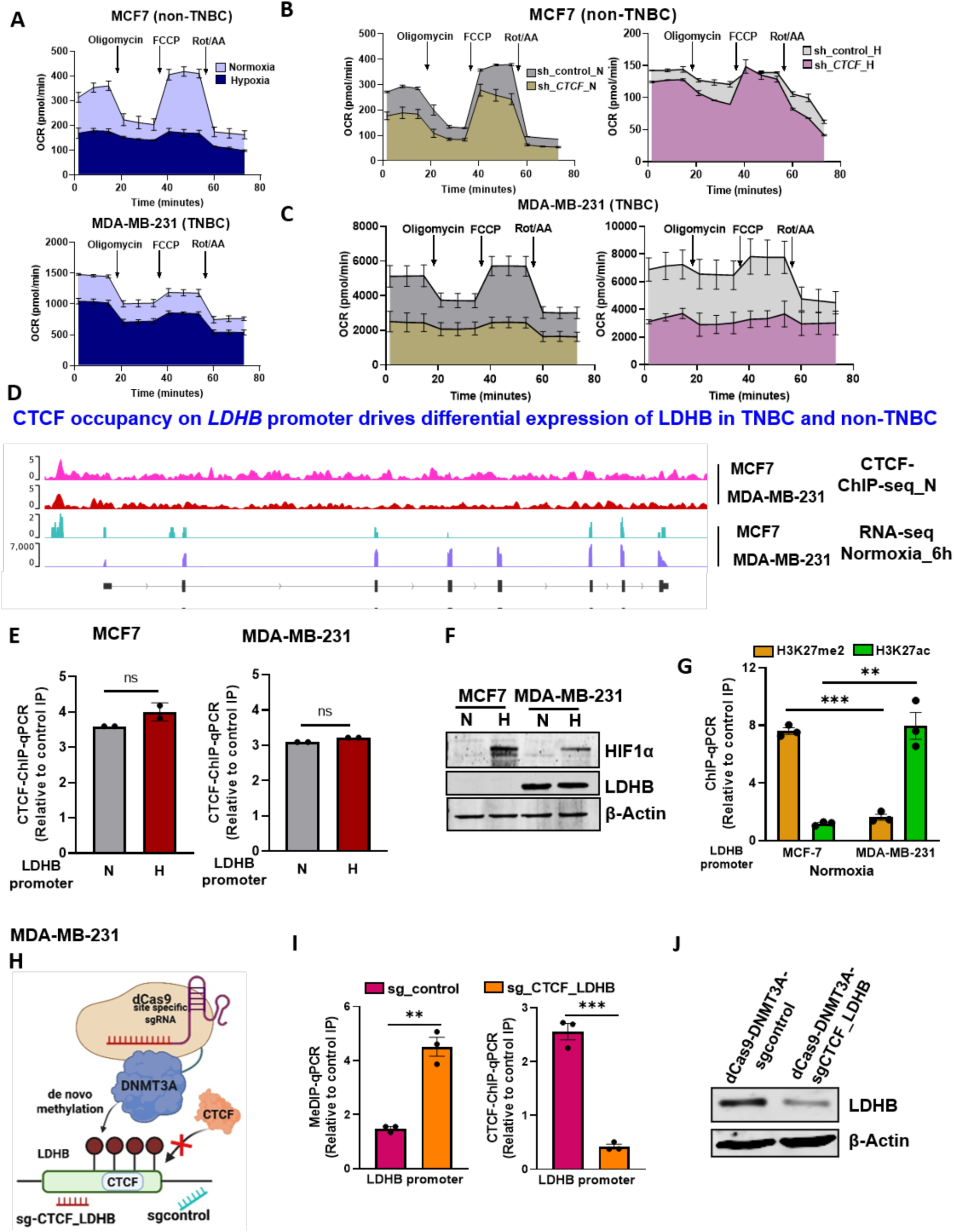
CTCF drives LDHB expression in TNBC. A) OCR analysis of MCF7 and MDA-MB-231 cells under normoxia and hypoxia B and C) OCR analysis of sh_control and sh_CTCF under hypoxia in B) MCF7 C) MDA-MB-231 cells. Cells were detected at basal level, followed by adding oligomycin (1.5µM), FCCP (0.5μM), and Rot/AA (0.5μM) D) genome browser view of LDHB loci of CTCF-ChIP-seq and RNA-seq under normoxia post 6 hours in MCF7 and MDA-MB-231 cells E) CTCF-ChIP-qPCR at LDHB promoter under normoxia and hypoxia in MCF7 and MDA-MB-231 cell lines F) Immunoblot of HIF1α and LDHB in normoxic and hypoxic MCF7 and MDA-MB-231 cells G) H3K27me2- and H3K27ac-ChIP-qPCR at LDHB promoter under normoxia and hypoxia in MCF7 and MDA-MB-231 cell lines H) Schematic representation of targeting the LDHB promoter by dCas9-DNMT3A with sgRNAs against CTCF to maintain methylation I and J) I) MeDIP-qPCR and CTCF-ChIP-qPCR on LDHB promoter and J) Immunoblot of LDHB in MDA-MB-231 cells transfected with dCas9-DNMT3A-sgCTCF versus dCas9-DNMT3A-sgControl. Data represent three technical replicates from pooled samples. Data are mean ± SEM, statistical significance was determined using unpaired two-tailed Student’s t-tests (E, G, I), ns non-significant, *p < 0.05, **p < 0.01, and ***p < 0.001; n = 3 biological replicates.

Because LDHB converts lactate to pyruvate and thereby feeds the TCA cycle ^14^ ^15^, its high expression offers an explanation for how MDA-MB-231 cells maintain OCR under hypoxia (**Figure 3A**). This raised the question of how CTCF, which occupies the *LDHB* promoter in both lines, can be associated with such different *LDHB* expression. We first addressed the role of CTCF in driving *LDHB* expression in MDA-MB-231 cells. To determine whether CTCF occupancy is necessary for *LDHB* expression in this TNBC line, we displaced CTCF using dCas9-DNMT3A with an sgRNA targeting the *LDHB* promoter (**Figure 3H**). De novo methylation displaced CTCF and reduced its occupancy (**Figure 3I**), and in turn reduced *LDHB* expression in MDA-MB-231 cells (**Figure 3J**). CTCF therefore drives *LDHB* expression in MDA-MB-231 cells (**Figure 3I**), accounting for the reduced OCR observed upon CTCF depletion under hypoxia (**Figure 3C**).

We next asked what CTCF occupancy at the LDHB promoter does in MCF7 cells, where *LDHB* is repressed. Interrogation of the LDHB promoter in the ChIP-Atlas database revealed ERα binding adjacent to the CTCF site in MCF7 cells (**Figure 4A**), which we validated by ChIP-qPCR in MCF7 cells (Figure 4B), consistent with MCF7 being ER-positive and MDA-MB-231 ER-negative. To dissect the mechanism governing differential *LDHB* expression in TNBC and non-TNBC cells, we performed luciferase reporter assays with an LDHB promoter construct. The LDHB reporter showed higher activity in MDA-MB-231 than in MCF7 cells (**Figure S2A**).

**Figure 4:**
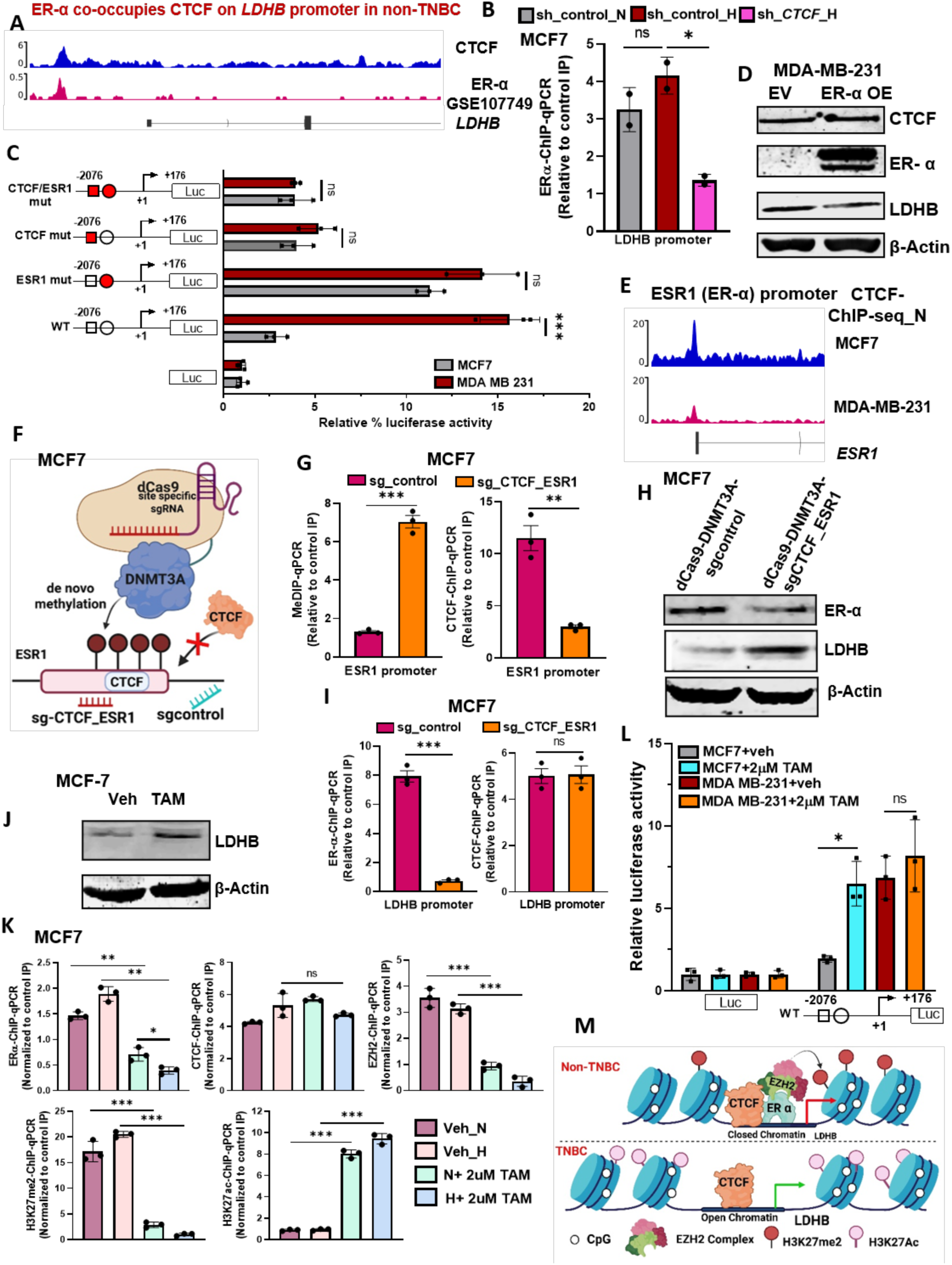
A CTCF-ERα complex represses LDHB in non-TNBC via modulating epigenetic landscape on LDHB promoter. A) genome browser view of LDHB loci of CTCF-ChIP-seq and ERα-ChIP-seq in MCF7 cells under normoxia B) ERα-ChIP-qPCR in CTCF depletion under hypoxia in comparison to sh_control under normoxic and hypoxic MCF7 cells C) MCF7 and MDA-MB-231 cells were co-transfected with wild type or CTCF or ERα or CTCF/ ERα mutant LDHB promoter luciferase construct (−337 to +115) and pRL-TK Renilla under normoxia D) Immunoblot of CTCF, ERα (to confirm overexpression of ERα) and LDHB in MDA-MB-231 cells with ectopically expressed either empty vector (EV) or ERα in MDA-MB-231 cells E) CTCF ChIP-seq signals at ESR1 promoter in MCF7 and MDA-MB-231 cells F) Schematic representation of targeting the ESR1 promoter by dCas9-DNMT3A with sgRNAs against CTCF to maintain methylation G and H) G) MeDIP-qPCR and CTCF-ChIP-qPCR on ESR1 promoter and H) Immunoblot of ERα and LDHB in MCF7 cells transfected with dCas9-DNMT3A-sgCTCF_ESR1 versus dCas9-DNMT3A-sgcontrol I) MeDIP-qPCR and CTCF-ChIP-qPCR on LDHB promoter in MCF7 cells transfected with dCas9-DNMT3A-sgCTCF_ESR1 versus dCas9-DNMT3A-sgcontrol J) Immunoblot of LDHB in MCF7 cells treated with Tamoxifen in comparison to the control cells under normoxia K) ERα, CTCF, EZH2, H3K27me2 and H3K27ac-ChIP-qPCR on LDHB promoter in MCF7 cells treated with Tamoxifen in comparison to the control cells under normoxia and hypoxia L) MCF7 and MDA-MB-231 cells were co-transfected with wild type LDHB promoter luciferase construct (−337 to +115) and pRL-TK Renilla and treated with Tamoxifen under normoxic condition M) Schematic showing CTCF-driven epigenetic and transcriptional regulation of LDHB expression in TNBC and non-TNBC breast cancer. Data are mean ± SEM, statistical significance was determined using one-way ANOVA with Tukey’s multiple comparisons test (B, C, L, M), unpaired two-tailed Student’s t-tests (G, J), ns non-significant, *p < 0.05, **p < 0.01, and ***p < 0.001; n = 3 biological replicates.

We then mutated the CTCF and ERα binding sites individually and together (double mutant) in the LDHB promoter construct (**Figure S2B**). Mutation of the ERα site markedly increased LDHB promoter activity in MCF7 cells with no effect in MDA-MB-231 cells (**Figure 4C**). Conversely, mutation of the CTCF motif alone did not change LDHB promoter activity in MCF7 cells but markedly reduced it in MDA-MB-231 cells, identifying the CTCF motif as a positive regulatory element in TNBC cells (**Figure 4C**). Mutation of both sites reduced LDHB promoter activity (**Figure 4C**). These data indicate that CTCF acts as a positive regulator of the LDHB promoter in TNBC, whereas in non-TNBC ERα co-occupies the promoter with CTCF and represses its expression. ERα binding at this site was dependent on CTCF occupancy, as ERα ChIP-qPCR in CTCF-depleted cells showed reduced ERα occupancy upon CTCF knockdown (**Figure 4B**).

To determine how the CTCF-ERα axis regulates *LDHB* expression in MCF7 cells, we overexpressed ERα and measured *LDHB* expression (**Figure 4D**). In addition to the LDHB promoter, we found CTCF occupancy at the *ESR1* promoter; this site was present in MCF7 but not in MDA-MB-231 cells and supports *ESR1* expression in MCF7 cells (**Figure 4E**). We therefore targeted dCas9-DNMT3A to the CTCF binding site at the ESR1 promoter (**Figure 4F**). This increased DNA methylation and reduced CTCF occupancy (**Figure 4G**), decreased ERα expression and increased *LDHB* expression in MCF7 cells (**Figure 4H**). ERα binding at the LDHB promoter was correspondingly reduced in dCas9-DNMT3A-sgCTCF cells relative to control cells (**Figure 4I**), while CTCF ChIP-qPCR showed that CTCF remained bound at the LDHB promoter (**Figure 4I**) and supported *LDHB* expression (**Figure 4H**).

These results indicate that CTCF is a positive driver of *LDHB* expression in breast cancer cells, and that the presence of ERα in MCF7 cells converts this input into repression. Consistent with this, ERα overexpression in MDA-MB-231 cells reduced *LDHB* expression (**Figure 4D**).

We next examined the epigenetic mechanism underlying *LDHB* repression in MCF7 cells. Treatment of MCF7 cells with tamoxifen (Tam), an ERα antagonist ^16^, increased *LDHB* expression (**Figure 4J**). Tam treatment reduced the occupancy of ERα and EZH2 and of the associated repressive H3K27me2 mark at the LDHB promoter without affecting CTCF binding, and increased H3K27ac at the same region (Figure 4K), thereby reconfiguring the MCF7 promoter towards the state observed in MDA-MB-231 cells. Consistently, luciferase assays in Tam-treated MCF7 cells showed higher LDHB promoter activity than controls (**Figure 4L**), and ERα depletion in MCF7 cells produced the same pattern (**Figure S2C and S2D**). Overall, these data indicate that CTCF supports *LDHB* expression in TNBC, whereas in MCF7 cells CTCF is accompanied by ERα, which assembles a repressive complex with EZH2 that deposits H3K27me2 ^17^ at the LDHB promoter and silences its expression (**Figure 4M**).

### LDHB expression determines the balance between H3K18 lactylation and acetylation under hypoxia

Lactate dehydrogenase B (LDHB) is a subunit of the lactate dehydrogenase complex that catalyses the reversible interconversion of lactate and pyruvate coupled to NAD+/NADH redox cycling and is thereby central to cellular metabolic balance ^14^. Beyond its established roles as an energy source and metabolic by-product, lactate is now recognized as a substrate for histone modification, placing metabolic reprogramming and epigenetic regulation, two hallmarks of cancer, in a direct reciprocal relationship ^6,18–20^.

Considering the above findings, we hypothesized that the differential expression of *LDHB* in MCF7 and MDA-MB-231 cells is associated with differential histone lactylation. To test this hypothesis, we measured pan-lactylation levels of histones in MCF7 cells overexpressing *LDHB* under hypoxia and observed that hypoxia increased pan-lactylation in MCF7 cells whereas *LDHB* overexpression abolished this induction under hypoxia (**Figure 5A**). To explore the relationship between *LDHB* and histone lactylation further, we assessed a panel of histone lactylation marks in MCF7 and MDA-MB-231 cells on transition from normoxia to hypoxia. H3K18la changed significantly in both cell lines, whereas H3K9la, H4K5la and H4K12la did not (**Figure 5B**). MCF7 cells showed higher H3K18la than MDA-MB-231 cells under normoxia, and under hypoxia H3K18la increased further in MCF7 cells while decreasing in MDA-MB-231 cells. We then examined *LDHB* expression and H3K18la at two time points under hypoxia in both lines (Figure 5C). Unexpectedly, MCF7 cells, which express little *LDHB*, showed higher H3K18la than MDA-MB-231 cells, which express *LDHB* (**Figure 5C**), indicating a strong inverse relationship between *LDHB* expression and H3K18la levels.

**Figure 5:**
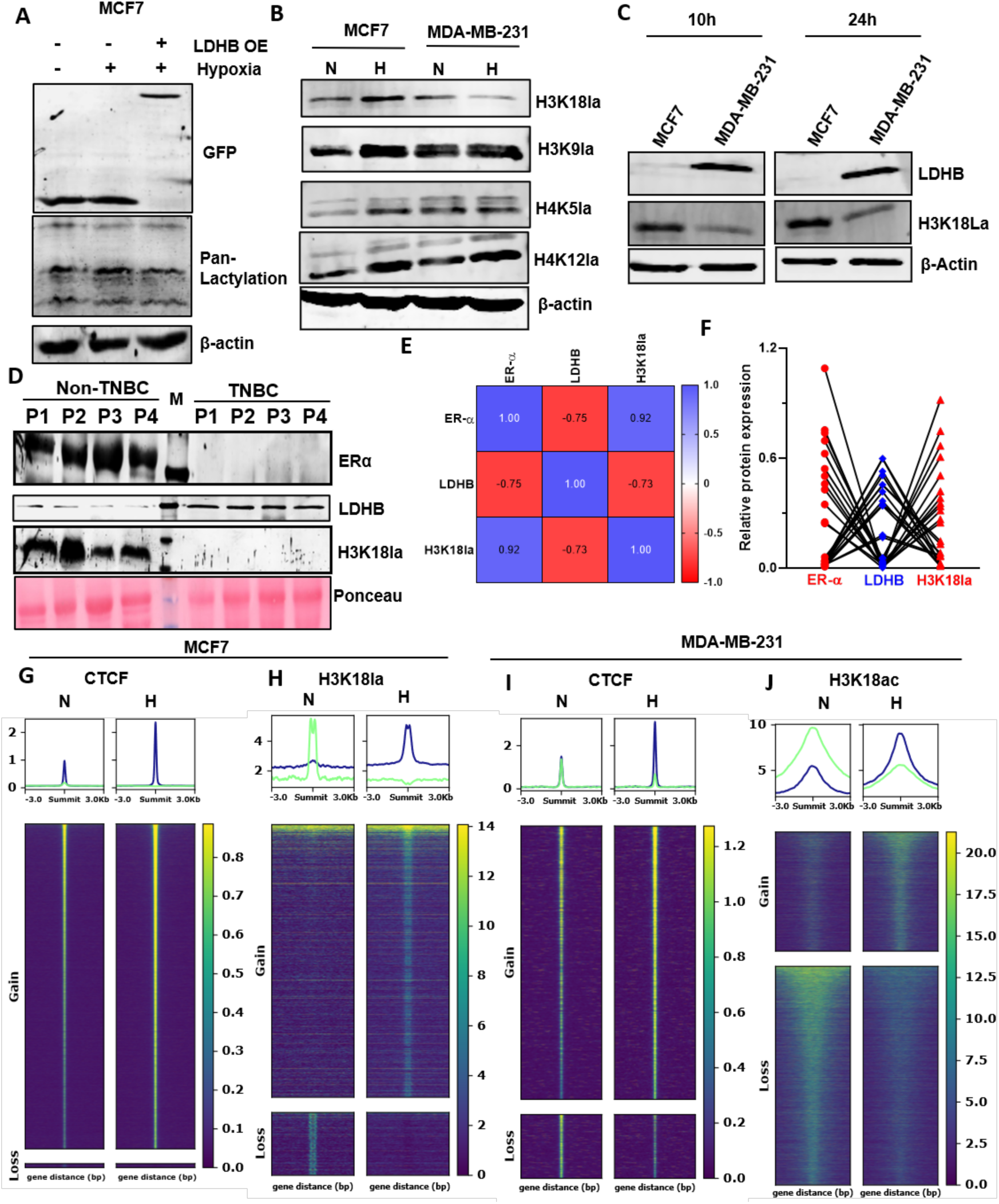
LDHB expression determines the balance between H3K18 lactylation and acetylation under hypoxia. A) Immunoblot of GFP (to confirm overexpression of LDHB) and Pan-lactylation with ectopically expressed either empty vector (EV) in normoxic and hypoxic or LDHB in hypoxic condition in MCF7 cells B) immunoblot of H3K18la, H3K9la, H4K5la and H4K12la in normoxic and hypoxic MCF7 and MDA-MB-231 cells C) immunoblot of H3K18la and LDHB at 10h and 24h in MCF7 and MDA-MB-231 cells post hypoxic treatment D) immunoblot of ERα, LDHB and H3K18la in breast tumor samples of non-TNBC and TNBC patients E and F) E) Pearson correlation plot and F) relative protein levels of ERα, LDHB and H3K18la in breast tumor samples of non-TNBC and TNBC patients (n=12 for each TNBC and non-TNBC samples) G-J) Heatmap of ChIP-seq signals of G) CTCF H) H3K18la in MCF7 I) CTCF and J) H3K18ac in MDA-MB-231 under normoxia and hypoxia post 10 hours. Peaks were grouped into gain and loss based on Differential binding analysis done via Diffbind.

To ask whether this relationship holds in human tumours, we examined the same pair of measurements in patient samples spanning both non-TNBC and TNBC (**Figure 5D and Figure S3A**). LDHB expression and H3K18la levels were strongly inversely correlated in patient samples (**Figure 5E and F**). Together, these findings indicate that global H3K18la levels are inversely linked to *LDHB* expression across breast cancer subtypes. To better understand this LDHB and its negative correlation with H3K18la in MCF7, we performed CTCF and H3K18la ChIP-seq while in MDA-MB-231, along with CTCF we performed H3K18ac (mutually exclusive competitive modification of H3K18la) under normoxia and hypoxia (**Figure 5G-J**). When comparing the ChIP-seq signals we observed significantly higher gained CTCF occupancy in both MCF7 and MDA-MB-231 corroborating with our previous study ^12^ (**Figure 5G and I, Table S1 and S2**). we also observed significantly higher gained H3K18la occupancy in MCF7 cells under hypoxia (**Figure 5J, Table S3**). Together, these analyses uncovered a strong association of H3K18la in LDHB negative MCF7 cells.

Lactate production is set by glycolytic rate, and hypoxia increases both ^21^. Both MCF7 and MDA-MB-231 cells showed increased intracellular and extracellular lactate on transition from normoxia to hypoxia (**Figure 6A**), and MDA-MB-231 cells had higher intracellular and extracellular lactate than MCF7 cells under both conditions (**Figure 6A**). This raises the question of how MDA-MB-231 cells show lower H3K18la than MCF7 cells despite their higher intracellular lactate.

**Figure 6:**
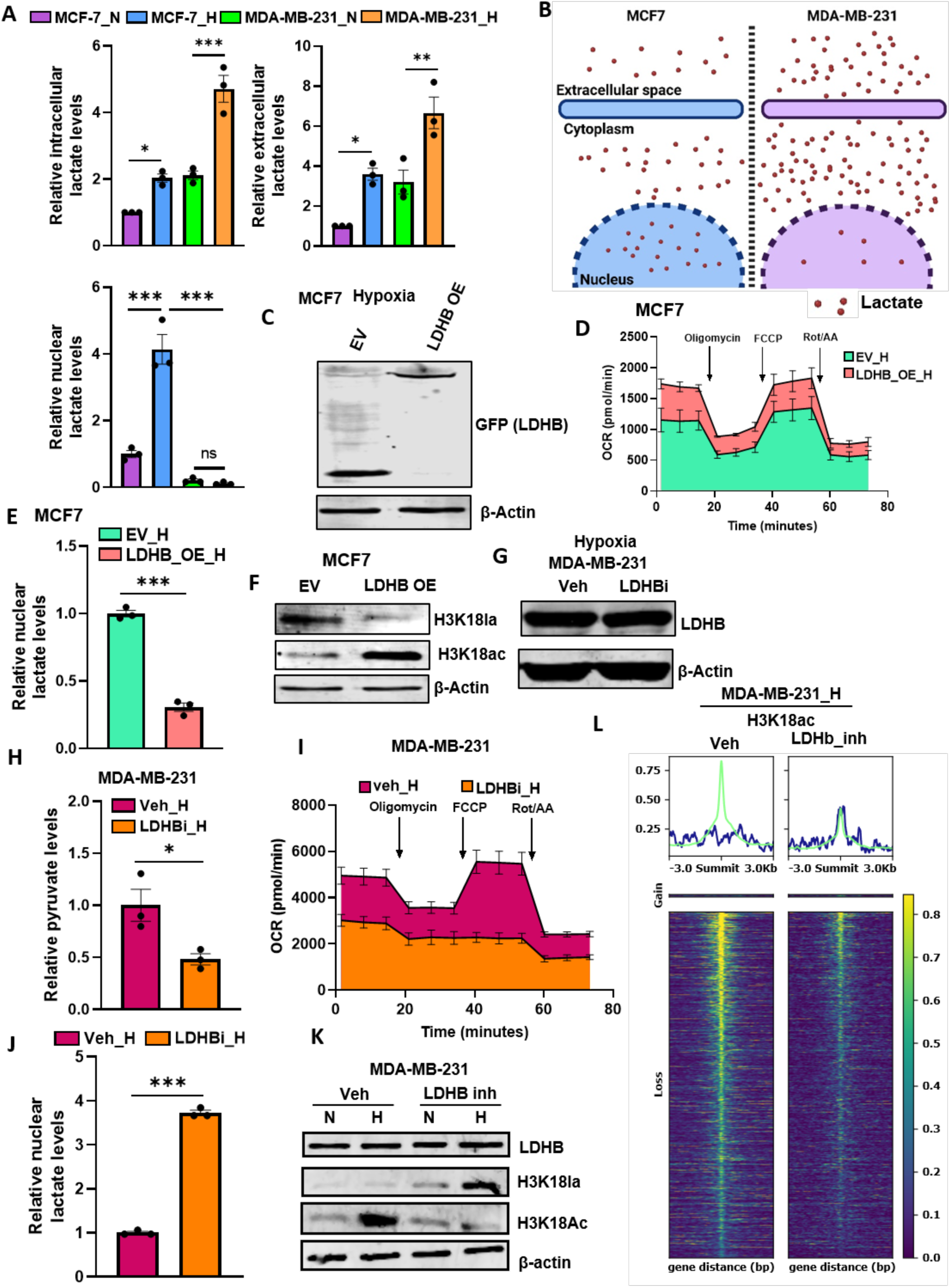
Nuclear lactate availability, rather than total intracellular lactate, drives H3K18 lactylation A) Intracellular, extracellular and nuclear lactate level in normoxic and hypoxic MCF7 and MDA-MB-231 cells B) schematic representing distribution of lactate across cytoplasm, extracellular space and nucleus in hypoxic MCF7 and MDA-MB-231 cells C-F) C) Immunoblot of GFP (to confirm overexpression of LDHB) D) OCR analysis E) Nuclear lactate levels F) immunoblot of H3K18la and H3K18ac in MCF7 cells with ectopically expressed either empty vector or LDHB in hypoxic condition G-L) G) Immunoblot of LDHB H) Pyruvate levels analysis I) OCR analysis J) Nuclear lactate levels K) immunoblot of LDHB, H3K18la and H3K18ac L) The heatmaps show the profile of H3K18ac in MDA-MB-231 cells treated with either vehicle (veh) or LDHB inhibitor (LDHBi) under hypoxia. Peaks were grouped into gain and loss based on Differential binding analysis done via Diffbind.Data are mean ± SEM, statistical significance was determined using one-way ANOVA with Tukey’s multiple comparisons test (A), unpaired two-tailed Student’s t-tests (E, H, J), ns non-significant, *p < 0.05, **p < 0.01, and ***p < 0.001; n = 3 biological replicates.

During glycolysis, glucose enters cells through glucose transporters (GLUTs) and is metabolized to pyruvate ^22^. Under baseline conditions pyruvate is largely imported into mitochondria and converted to acetyl-CoA for the tricarboxylic acid (TCA) cycle, while a fraction is reduced to lactate by lactate dehydrogenase in the cytosol, maintaining an equilibrium between oxidative phosphorylation and aerobic glycolysis ^23^. Lactate can be shuttled across membranes by monocarboxylate transporters (MCT1/MCT4) ^24^, and is converted to lactyl-CoA by acyl-coenzyme A (acyl-CoA) synthetase short-chain family member 2 (ACSS2); lactyl-CoA is then transferred to lysine residues on histone and non-histone proteins by acetyltransferases including E1A binding protein p300 (p300) and CREB-binding protein (CBP), linking metabolic state to epigenetic regulation ^7^. Under hypoxic or hyper glycolytic conditions, nuclear lactate accumulation drives p300-mediated lactylation, converting metabolic signals into epigenetic instructions ^7^.

Because histone lactylation is templated on the nuclear lactate pool rather than on total cellular lactate, nuclear lactate concentration is the more proximal determinant of the extent of histone lactylation. In addition, high lactate has recently been reported to be sensed by AARS1, which can lactylate proteins including histone H3 at K18 ^25,26^.

We therefore measured nuclear lactate in MCF7 and MDA-MB-231 cells under normoxia and hypoxia. Despite their higher intracellular lactate, MDA-MB-231 cells contained markedly lower free nuclear lactate, whereas MCF7 cells contained free nuclear lactate that increased further under hypoxia (**Figure 6A and 6B**).

These findings indicate that nuclear, rather than total intracellular, lactate availability sets the level of H3K18 lactylation. We next asked whether nuclear lactate is related to LDHB, given the inverse relationship between LDHB and H3K18la. We overexpressed *LDHB* in MCF7 cells and maintained them under hypoxia (**Figure 6C**). *LDHB* overexpression increased OCR relative to control (**Figure 6D**), consistent with lactate being converted to pyruvate and directed into oxidative phosphorylation. Nuclear lactate was correspondingly reduced in *LDHB*-overexpressing cells under hypoxia (**Figure 6E**), as was H3K18la (**Figure 6F**). Because H3K18 lactylation and acetylation are mutually exclusive at the same residue, and because *LDHB* overexpression is expected to increase acetyl-CoA availability by channelling lactate-derived pyruvate into the TCA cycle, we also measured H3K18ac and found it increased in LDHB-overexpressing cells relative to control cells under hypoxia (**Figure 6F**). To test the converse, we treated MDA-MB-231 cells with the LDHB inhibitor AXKO-0046 (LDHBi; MedChemExpress) under hypoxia (**Figure 6G**). LDHBi reduced pyruvate levels (**Figure 6H**) and reduced OCR (**Figure 6I**), consistent with inhibition of lactate-to-pyruvate conversion.

LDHBi increased nuclear lactate relative to control cells under hypoxia (**Figure 6J**), reduced H3K18ac and increased H3K18la (**Figure 6K**). Genome-wide, H3K18ac ChIP-seq showed globally reduced H3K18ac in MDA-MB-231 cells treated with LDHBi under hypoxia (**Figure 6L, Table S4**).

Thus hypoxia-induced lactate accumulation raises nuclear lactate and H3K18la in MCF7 cells, and LDHB overexpression reverses this; conversely, LDHB inhibition in MDA-MB-231 cells raises nuclear lactate. Together these data identify LDHB as a determinant of H3K18 lactylation in breast cancer cells under hypoxia and show that LDHB expression is directly associated with H3K18ac levels under the same conditions. Overall, our data indicate that nuclear lactate availability, rather than total intracellular lactate, drives H3K18 lactylation, converting metabolic signals into epigenetic instructions.

### CTCF regulates MCT1 expression, and CTCF-dependent MCT1 nuclear localization sustains high nuclear lactate under hypoxia in non-TNBC

This observation raises two linked questions: how MDA-MB-231 cells maintain low nuclear lactate despite high intracellular lactate, and what delivers lactate into the nucleus of MCF7 cells to sustain a gradient sufficient for incorporation at H3K18.

We first observed that CTCF depletion in MCF7 cells under hypoxia reduced H3K18la levels (**Figure 7A**) and reduced nuclear lactate levels (**Figure 7B**). We therefore hypothesized that CTCF contributes not only to glycolysis but also to nuclear lactate delivery under hypoxia.

**Figure 7:**
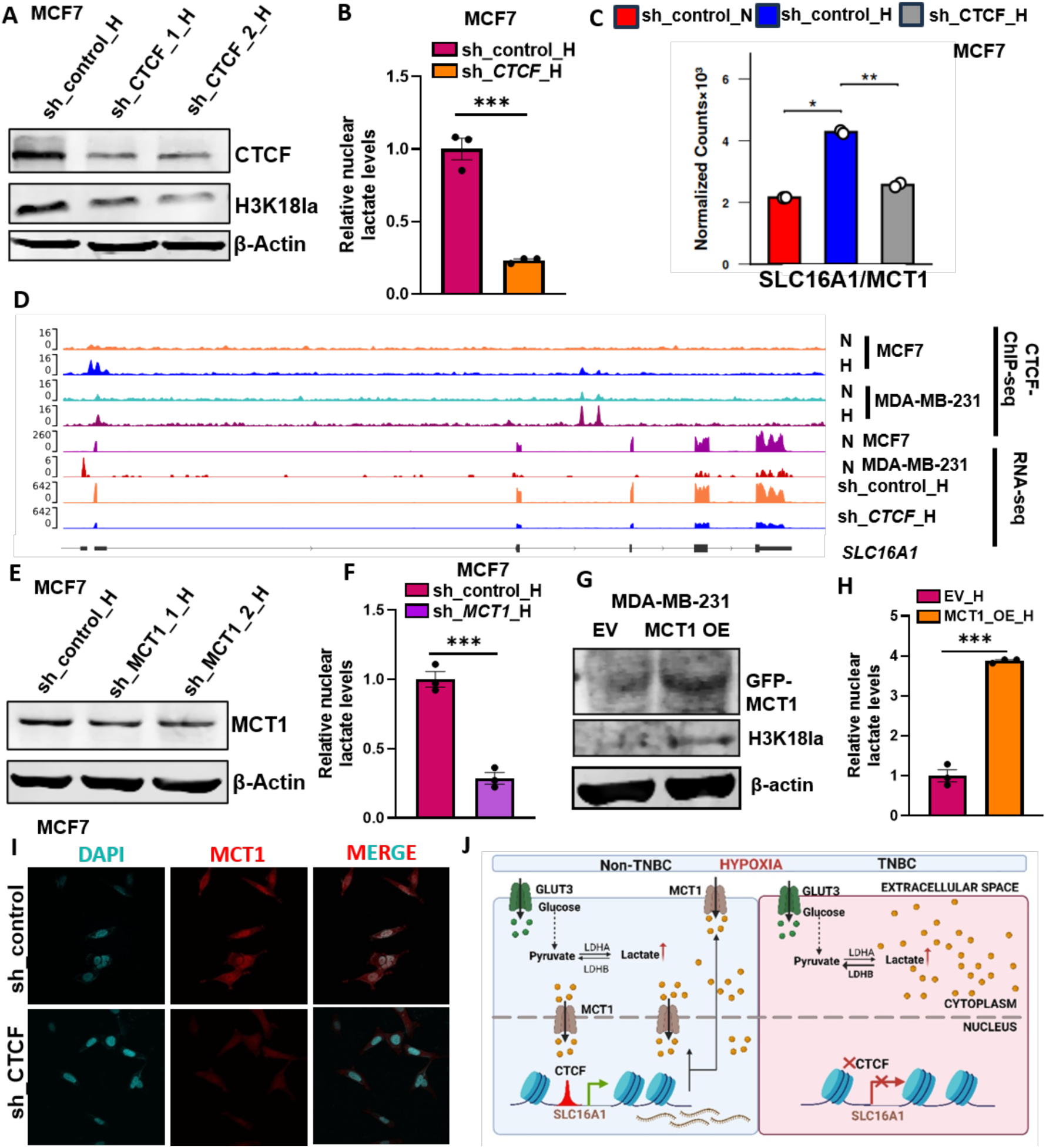
CTCF regulates MCT1 expression, and CTCF-dependent MCT1 nuclear localization sustains high nuclear lactate under hypoxia in non-TNBC. A-B) A) Immunoblot of CTCF and H3K18la B) nuclear lactate levels in CTCF knockdown and shControl MCF7 cells under hypoxia C) mRNA expression of SLC16A1 in sh_control under normoxia and hypoxia and sh_CTCF under hypoxia in MCF7 cells inferred from CTCF depletion RNA-seq D) genome browser view of SLC16A1/MCT1 loci of CTCF-ChIP-seq in MCF7 and MDA-MB-231 under normoxia and hypoxia, RNA-seq under normoxia post 6 hours in MCF7 and MDA-MB-231 cells and RNA-seq in shcontrol and shCTCF under hypoxia in MCF7 cells E-F) E) immunoblot of MCT1 F) nuclear lactate levels in MCT1 knockdown and shControl MCF7 cells under hypoxia G-H) G) immunoblot of Immunoblot of eGFP (to confirm overexpression of MCT1) and H3K18la H) nuclear lactate levels in MDA-MB-231 cells with ectopically expressed either empty vector or MCT1 in hypoxic condition I) Schematic showing CTCF-driven transcriptional regulation of MCT1 expression in non-TNBC and TNBC breast cancer. Data are mean ± SEM, statistical significance was determined using unpaired, two-tailed Welch’s Two-Sample t-test (C), unpaired two-tailed Student’s t-tests (B, F, H), ns non-significant, *p < 0.05, **p < 0.01, and ***p < 0.001; n = 3 biological replicates.

To identify the responsible transporter, we examined the RNA-seq data from CTCF-knockdown MCF7 cells under hypoxia for genes involved in lactate transport that were downregulated upon CTCF knockdown. CTCF knockdown significantly altered several genes associated with protein and macromolecular transport, among which *SLC16A1* (MCT1) was downregulated upon CTCF knockdown in MCF7 cells under hypoxia (**Figure 7C, Table S1**).

MCT1 is a lactate transporter that imports lactate into cells ^24^. CTCF occupied the MCT1 promoter in MCF7 cells, and this occupancy was absent in MDA-MB-231 cells (**Figure 7D, Table S1 and S2**). Our RNA-seq data showed that *MCT1* expression increased under hypoxia relative to normoxia and decreased upon CTCF knockdown under hypoxia (**Figure 7D**). To test the contribution of MCT1 to nuclear lactate, we knocked down *MCT1* in MCF7 cells (**Figure 7E**) and observed reduced nuclear lactate levels (**Figure 7F**). Conversely, overexpression of *MCT1* in MDA-MB-231 cells under hypoxia (**Figure 7G**) increased both nuclear lactate and H3K18la relative to control cells (**Figure 7G and H**). Finally, because MCT1 at the nuclear membrane would be expected to govern nuclear lactate levels, we used confocal imaging to examine MCT1 localization. Under hypoxia, *MCT1* expression increased and its localization to the nuclear membrane also increased relative to normoxic control (**Figure 7I**). CTCF knockdown reduced both *MCT1* expression and its nuclear membrane localization, consistent with the reduced nuclear lactate levels observed in CTCF-depleted cells under hypoxia (**Figure 7I**). overall, we conclude that CTCF regulates *MCT1* expression in MCF7 cells, and CTCF-dependent MCT1 nuclear localization sustains high nuclear lactate under hypoxia in non-TNBC (**Figure J**).

### LDHB expression is positively associated with MCT4 expression and high ECAR under hypoxia in TNBC

LDHB expression is inversely associated with nuclear lactate and H3K18la, while its presence promotes H3K18ac in MDA-MB-231 cells under hypoxia. We therefore asked how LDHB-associated H3K18ac shapes gene expression in TNBC, which is more aggressive and harder to treat than non-TNBC. We performed RNA-seq in MDA-MB-231 cells treated with LDHBi under hypoxia. A substantial number of genes were significantly downregulated upon *LDHB* inhibition under hypoxia (**Figure 8A, Table S4**), and these were enriched for hypoxia, glycolysis, oxidative phosphorylation and other hallmark pathways (**Figure 8B, Table S4**). Intersecting the promoters of the downregulated genes with H3K18ac ChIP-seq from LDHBi-treated cells showed that 595 downregulated genes had reduced promoter H3K18ac relative to control under hypoxia (**Figure 8C, Table S4**), among which *SLC16A3* (MCT4) was downregulated upon LDHB inhibitor treatment in MDA-MB-231 cells under hypoxia (**Figure 8D, Table S4**).

**Figure 8:**
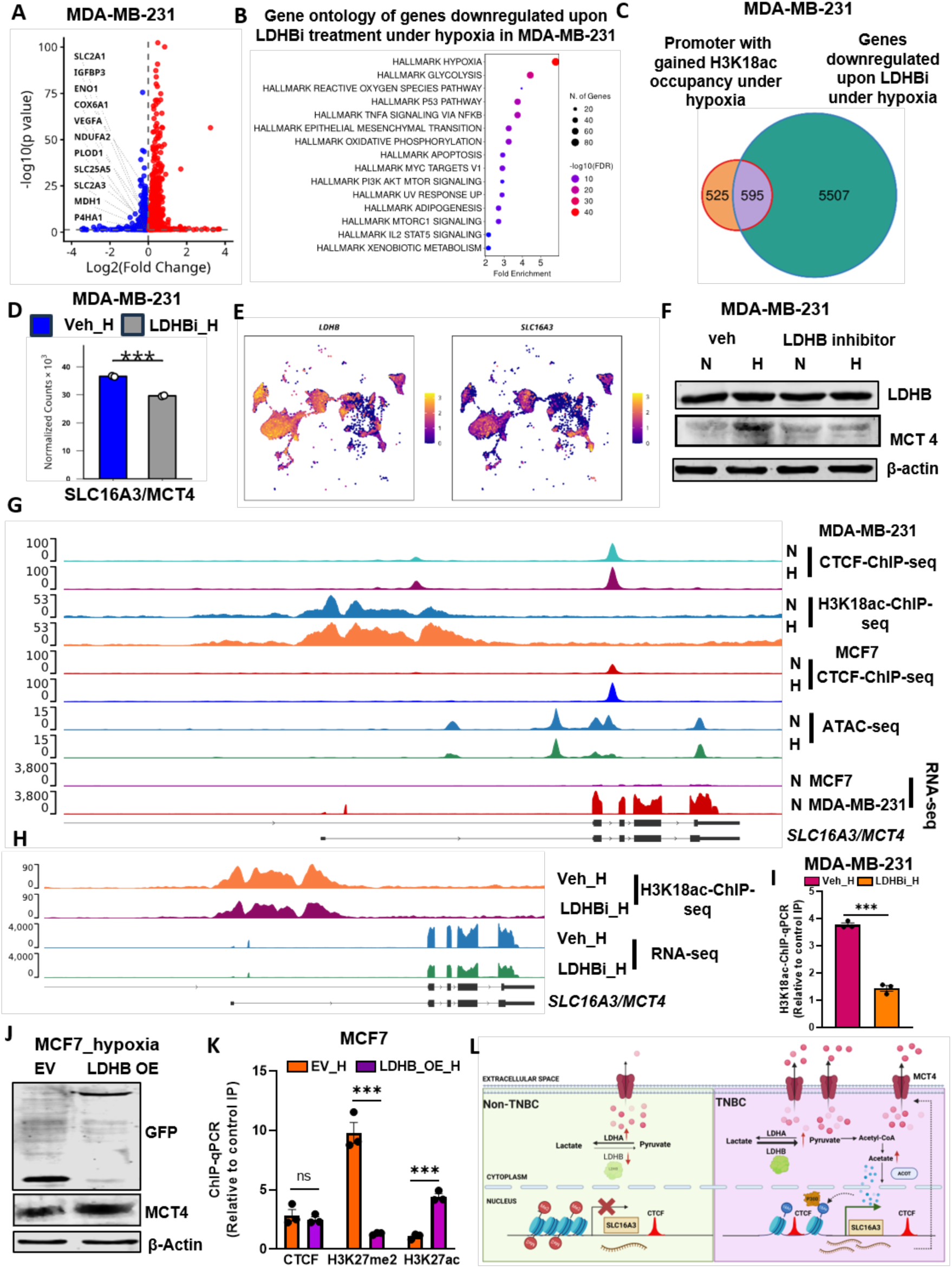
LDHB expression is positively associated with MCT4 expression and high ECAR under hypoxia in TNBC. A-B) A) Volcano plot of differentially expressed genes B) Representative significantly enriched terms from the Gene Ontology of genes downregulated upon vehicle control and LDHB inhibitor treatment in MDA-MB-231 cells under hypoxia C) Venn diagram showing overlap of downregulated genes with reduced promoter H3K18ac in LDHBi relative to control under hypoxia D) mRNA expression of SLC16A3 in inferred from LDHBi RNA-seq in MDA-MB-231 E) UMAP plot of breast epithelial cancerous cell populations stratified as continuous hypoxic state using Buffa score highlighting LDHB expression and SLC16A3 expression F) Immunoblot of LDHB and MCT4 in MDA-MB-231 cells treated with either vehicle (veh) or LDHB inhibitor (LDHBi) under normoxia and hypoxia G) genome browser view of SLC16A3/MCT4 loci of CTCF- and H3K18la-ChIP-seq in MDA-MB-231 under normoxia and hypoxia, CTCF-ChIP-seq, ATAC-seq in MCF7 under normoxia and hypoxia and RNA-seq under normoxia post 6 hours in MCF7 and MDA-MB-231 cells H) genome browser view of H3K18ac-ChIP at MCT4 promoter and RNA-seq for MCT4 in MDA-MB-231 cells treated with either vehicle (veh) or LDHB inhibitor (LDHBi) under hypoxia I) ChIP-qPCR of H3K18ac at MCT4 promoter in MDA-MB-231 cells treated with either vehicle (veh) or LDHB inhibitor (LDHBi) under hypoxia J-K) J) Immunoblot of GFP (to confirm overexpression of LDHB) and MCT4 K) ChIP-qPCR of H3K27me2 and H3K27ac at CTCF occupancy loci in the 1st intron of MCT4 in MCF7 cells with ectopically expressed either empty vector or LDHB in hypoxic condition L) Schematic showing LDHB associated H3K18ac mark and CTCF-driven epigenetic and transcriptional regulation of SLC16A3 expression in TNBC and non-TNBC. Data are mean ± SEM, statistical significance for Co-expression was assessed using Fisher’s exact test (odds ratio = 1.75, p < 2.2 × 10⁻¹⁶) (E), unpaired two-tailed Student’s t-tests (J), ns non-significant, *p < 0.05, **p < 0.01, and ***p < 0.001; n = 3 biological replicates.

Further, we examined the relationship between LDHB and MCT4 more directly. Analysis of scRNA-seq data showed a positive co-expression of LDHB and MCT4 in malignant cells (Figure 8E). *MCT4* expression increased under hypoxia and this increase was blunted by LDHBi in MDA-MB-231 cells (**Figure 8F**). Integrating CTCF and H3K18ac ChIP-seq with RNA-seq in normoxic MCF7 and MDA-MB-231 cells showed that CTCF occupies the MCT4 promoter in MDA-MB-231 but not in MCF7 cells, and that *MCT4* expression is higher in MDA-MB-231 than MCF7 cells (**Figure 8G, Table S1-4**). We also found CTCF occupancy within intron 1 in both MCF7 and MDA-MB-231 cells, accompanied by reduced ATAC-seq signal at this site in MCF7 under hypoxia (**Figure 8G**).

Furthermore, analysis of H3K18ac ChIP-seq and RNA-seq in LDHB inhibitor-treated MDA-MB-231 cells showed reduced H3K18ac occupancy at the *SLC16A3* (MCT4) promoter and reduced MCT4 expression under hypoxia (**Figure 8H**). H3K18ac ChIP-qPCR at the MCT4 promoter confirmed reduced occupancy upon LDHBi treatment under hypoxia (**Figure 8I**).

Overexpression of LDHB in MCF7 cells increased *MCT4* expression (**Figure 8J**) and was accompanied by reduced H3K27me2 and increased H3K27ac at the locus, while CTCF occupancy remained unchanged (**Figure 8K**). Together these data suggest that CTCF occupancy at intron 1 represses *MCT4* in MCF7 cells, whereas in MDA-MB-231 cells CTCF occupancy together with LDHB-dependent H3K18ac activates *MCT4* expression, contributing to the higher ECAR of these cells (**Figure 8L**).

### Coupling of CTCF-mediated Pol II pausing with an H3K18la/ac adaptor mechanism directs divergent splicing outcomes and drives stemness in TNBC and EMT in non-TNBC under hypoxia

Given the genome-wide association between CTCF, H3K18la/ac and splicing (**Figure S4, Table S5**), we next assessed the biological consequences of these splicing events, focusing on those with divergent outcomes in TNBC and non-TNBC cancer cells. We identified a splicing event at exon 3 of *TACC3* (**Figure 9A**) whose outcome differed between TNBC and non-TNBC cells (**Figure 9B and Figure S5A**). We pursued this event and first validated the RNA-seq findings across a panel of TNBC and non-TNBC cell lines. TNBC lines consistently included exon 3 (full-length, FL) under hypoxia, whereas non-TNBC lines skipped exon 3 (shorter isoform, Δ3) under hypoxia (**Figure 9C, Table S5**). Thus, TNBC cells favour exon 3 inclusion while non-TNBC cells favour its exclusion from mature TACC3 mRNA under hypoxia. The same pattern was evident in TCGA data (TSVdb), where TNBC tumours showed higher exon 3 inclusion in TACC3 mRNA (**Figure 9D**), and the FL isoform also increased under hypoxia at the protein level (**Figure 9E**).

**Figure 9:**
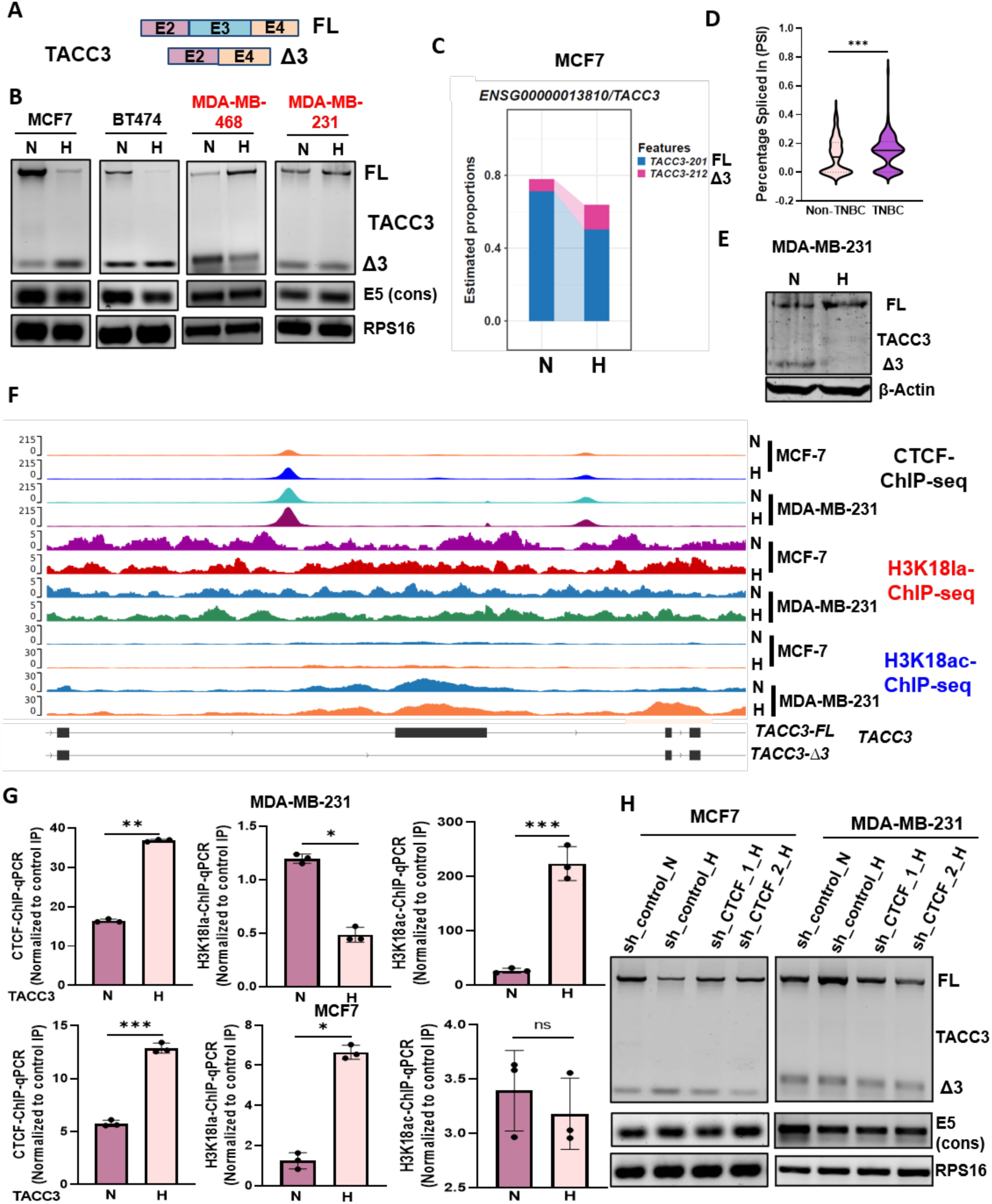
CTCF-mediated TACC3 splicing outcome in breast cancer subtypes under hypoxia. A) schematic representing TACC3 gene alternative splicing, exon3 inclusion give rise to TACC3-full length (FL) and exclusion give rise to Δ3 isoform B) alternative splicing analysis of TACC3 exon 3 in various breast cancer cell lines (non-TNBC and TNBC) under normoxia and hypoxia C) isoform level quantification of TACC3-ENST00000313288 (FL) and TACC3-ENST00000612220 (Δ3 isoform) using SALMON inferred from RNA-seq performed in MCF7 under normoxia and hypoxia D) percentage spliced In (PSI) of FL isoform in TCGA database stratified as TNBC and Non-TNBC E) immunoblot of TACC3 in MDA-MB-231 cells under normoxia and hypoxia showing both FL and Δ3 isoform F) genome browser view of TACC3 exon 3 loci of CTCF-, H3K18la-and H3K18lac-ChIP-seq in MCF7 and MDA-MB-231 under normoxia and hypoxia post 10 hours G) CTCF-, H3K18la- and H3K18ac-ChIP-qPCR at CBSDS loci of TACC3 gene region in MCF7 and MDA-MB-231 cells under normoxia and hypoxia H) alternative splicing analysis of TACC3 exon 3 in CTCF knockdown and shControl in MCF7 and MDA-MB-231 cells under hypoxia. Data are mean ± SEM, statistical significance was determined using Wilcoxon signed rank test (D), unpaired two-tailed Student’s t-tests (G), ns non-significant, *p < 0.05, **p < 0.01, and ***p < 0.001; n = 3 biological replicates.

This observation prompted us to define the molecular mechanism driving the divergent splicing outcome in TNBC and non-TNBC cells, and its consequences for cancer progression under hypoxia.

We first analysed CTCF occupancy around the spliced region in both cell lines and found three CTCF-occupied sites associated with exon 3: (1) a site 1270 bp upstream (CBSUS); (2) a site overlapping exon 3 (CBSO); and (3) a site 1128 bp downstream of exon 3 (CBSDS) (**Figure 9F, Table S1-3**). CBSUS and CBSDS were present in both MCF7 and MDA-MB-231 cells, whereas CBSO was detected only in MCF7 cells (**Figure 9F**). H3K18la ChIP-seq showed increased H3K18la around CBSDS under hypoxia only in MCF7 cells, while H3K18ac ChIP-seq showed increased H3K18ac around CBSDS under hypoxia only in MDA-MB-231 cells (**Figure 9F**). CTCF occupancy at CBSDS is therefore associated with H3K18la in MCF7 cells and with H3K18ac in MDA-MB-231 cells under hypoxia (**Figure 9F**). We validated these observations by ChIP-qPCR, confirming increased CTCF and H3K18la occupancy in MCF7 cells and increased CTCF and H3K18ac occupancy in MDA-MB-231 cells at CBSDS under hypoxia (**Figure 9G**).

We next examined the splicing outcome in both cell lines following CTCF depletion under hypoxia. In MCF7 cells, CTCF depletion reduced the Δ3 isoform and increased the FL isoform under hypoxia (**Figure 9H**). In MDA-MB-231 cells, CTCF depletion had the opposite effect, increasing Δ3 and reducing FL (**Figure 9H**). CTCF is therefore required for the TACC3 splicing outcome in both subtypes, but the direction of its effect differs between TNBC and non-TNBC cell lines under hypoxia.

### Targeted de novo methylation of the CTCF binding site at CBSDS alters CTCF-mediated TACC3 exon 3 alternative splicing under hypoxia

CTCF-mediated Pol II pausing is known to influence splicing outcome ^12,27^. To test whether CTCF-mediated RNA Pol II pausing is sufficient to direct TACC3 splicing under hypoxia, we targeted the dCas9-DNMT3A system with an sgRNA against CBSDS downstream of TACC3 exon 3 (**Figure 10A**). MeDIP-qPCR confirmed increased methylation at CBSDS in dCas9-DNMT3A-sgCTCF as compared to dCas9-DNMT3A-sgcontrol (**Figure 10B**), and CTCF occupancy was correspondingly reduced (**Figure 10B**). Semi-quantitative PCR and qPCR showed a significant switch in the splicing outcome of exon 3, with reduced Δ3 isoform and increased FL isoform in dCas9-DNMT3A-sgCTCF as compared to dCas9-DNMT3A-sgcontrol under hypoxia (**Figure 10C and D**). H3K18la and RNA Pol II occupancy at CBSDS were also reduced in dCas9-DNMT3A-sgCTCF cells as compared to dCas9-DNMT3A-sgcontrol under hypoxia (**Figure 10E**). These data indicate that hypoxia-driven, CTCF-mediated RNA Pol II pausing is sufficient to direct alternative splicing of TACC3 exon 3 towards exclusion under hypoxia. The concomitant loss of H3K18la at CBSDS alongside loss of exon 3 exclusion prompted us to examine the CTCF-H3K18la relationship more closely.

**Figure 10:**
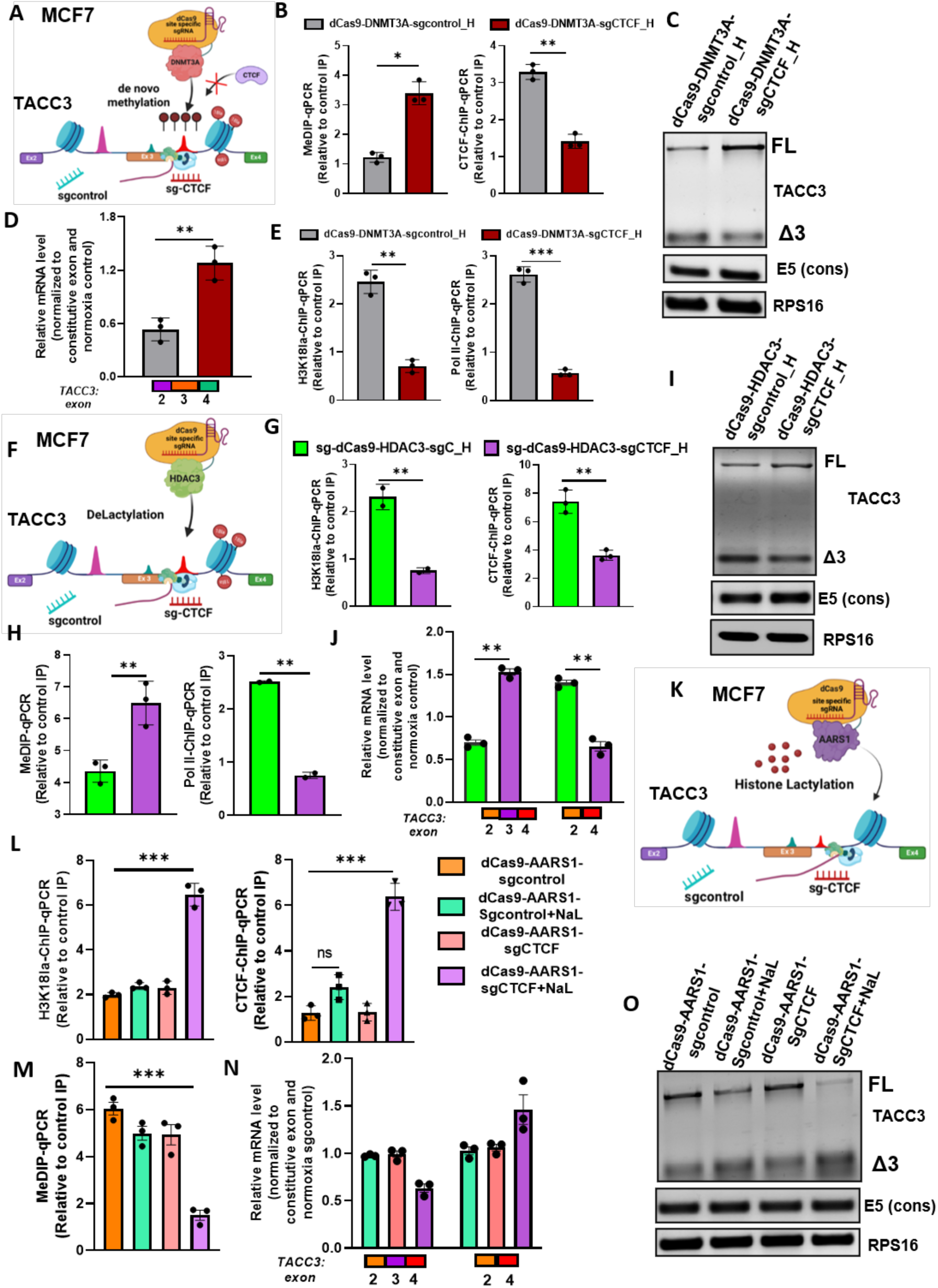
CTCF and H3K18la together direct TACC3 splicing towards exon 3 exclusion in the non-TNBC. A) Schematic representation of targeting the CTCF motif at CBSDS loci of TACC3 gene region by dCas9-DNMT3A with specific sgRNA against CTCF to maintain methylation and suppress CTCF occupancy B) MeDIP-qPCR and CTCF-ChIP-qPCR at CBSDS loci of TACC3 gene region (C) alternative splicing analysis of TACC3 D) qPCR analysis of alternative splicing of TACC3 E) H3K18la and Pol II-ChIP-qPCR at CBSDS loci of TACC3 gene region in MCF7 cells transfected with dCas9-DNMT3A-sgCTCF in comparison to the sgcontrol under hypoxia F) Schematic representation of targeting the CTCF motif at CBSDS loci of TACC3 gene region by dCas9-HDAC3 with specific sgRNA used in (A) to remove H3K18la G) MeDIP-qPCR and CTCF-ChIP-qPCR at CBSDS loci of TACC3 gene region (H) H3K18la and Pol II-ChIP-qPCR I) alternative splicing analysis of TACC3 J) qPCR analysis of alternative splicing of TACC3 at CBSDS loci of TACC3 gene region in MCF7 cells transfected with dCas9-HDAC3-sgCTCF in comparison to the sgcontrol under hypoxia K) Schematic representation of targeting the CTCF motif at CBSDS loci of TACC3 gene region by dCas9-AARS1 with specific sgRNA used in (A) to add H3K18la L) H3K18la-and CTCF-ChIP-qPCR (M) MeDIP-qPCR at CBSDS loci of TACC3 gene region N) qPCR analysis of alternative splicing of TACC3 O) alternative splicing analysis of TACC3 in MCF7 cells transfected with dCas9-AARS1-sgCTCF in comparison to the sgcontrol either supplemented solvent or with 20mm NaLa under normoxia. Data are mean ± SEM, statistical significance was determined using unpaired two-tailed Student’s t-tests (B, D, E, G, H, J), one-way ANOVA with Tukey’s multiple comparisons test (L, M, N), ns non-significant, *p < 0.05, **p < 0.01, and ***p < 0.001; n = 3 biological replicates.

### Targeted removal of H3K18la at CBSDS alters *TACC3* exon 3 alternative splicing under hypoxia

H3K18la at CBSDS was reduced when CTCF occupancy was abolished by targeted methylation of CBSDS. We therefore tested the role of H3K18la in TACC3 exon 3 splicing directly, using a dCas9-HDAC3 construct to remove H3K18la at this site (HDAC3 has been reported as an eraser of H3K18la ^28^ (**Figure 10F**). dCas9-HDAC3-sgCTCF reduced H3K18la at CBSDS as compared to dCas9-HDAC3-sgcontrol under hypoxia (**Figure 10G**). CTCF and Pol II occupancy were also reduced, and DNA methylation increased, in dCas9-HDAC3-sgCTCF as compared to dCas9-HDAC3-sgcontrol under hypoxia (**Figure 10G and H**). Correspondingly, the FL isoform increased and the Δ3 isoform decreased in dCas9-HDAC3-sgCTCF as compared to dCas9-HDAC3-sgcontrol under hypoxia (**Figure 10I and J**). Together, these observations indicate that CTCF-mediated Pol II pausing and H3K18la at CBSDS act in concert to direct TACC3 exon 3 splicing under hypoxia.

### Targeted de novo lactylation of H3K18 at CBSDS remodels the local epigenetic landscape and directs TACC3 exon 3 alternative splicing under normoxia

To define the interdependence between H3K18la and CTCF occupancy at CBSDS of TACC3 exon 3, we remodelled the epigenetic landscape of normoxic cells. AARS1 has recently been reported as a specific writer of H3K18la under high lactate concentrations ^25,26^. We therefore constructed a dCas9-AARS1 system and catalysed targeted de novo lactylation of H3K18 at CBSDS under normoxia (**Figure 10K**). Cells transfected with dCas9-AARS1-sgcontrol, with or without 20 mM sodium lactate (NaL) supplementation, showed no induction of H3K18la at the CBSDS locus (**Figure 10L**). dCas9-AARS1-sgCTCF alone also failed to induce H3K18la, but in the presence of NaL it increased H3K18la at CBSDS (**Figure 10L**). Under these conditions we observed increased CTCF occupancy and decreased DNA methylation at CBSDS in dCas9-AARS1-sgCTCF+NaL as compared to the other controls under normoxia (**Figure 10L and M**), together with a significant decrease in the TACC3 FL isoform (**Figure 10N and O**). Taken together with the preceding results, these data indicate that CTCF and H3K18la together direct TACC3 splicing towards exon 3 exclusion in the non-TNBC cell line. The observation that induced H3K18la at CBSDS also decreased local DNA methylation suggests a further link between H3K18la and DNA methylation that will require dedicated study.

### LDHB is positively associated with TACC3 exon 3 inclusion under hypoxia in TNBC cells

Based on the observation that LDHB has a direct impact on H3K18 lactylation status under hypoxia, we next asked whether LDHB expression is associated with the TACC3 splicing outcome. Overexpression of LDHB in MCF7 cells increased the FL isoform relative to empty vector under hypoxia, consistently across RNA-seq, qPCR and semi-quantitative PCR (**Figure 11A and B**).

**Figure 11:**
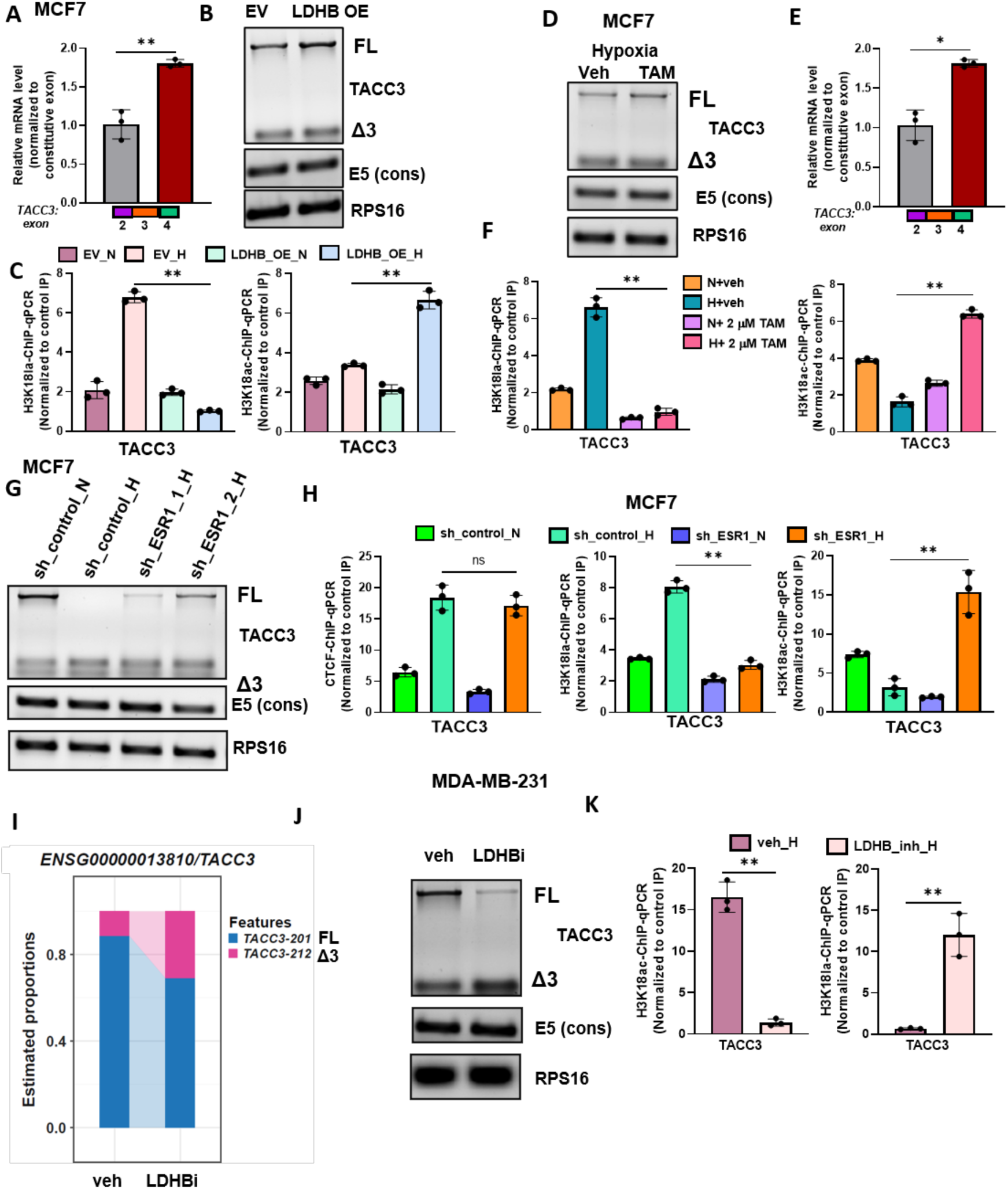
LDHB is positively associated with TACC3 exon 3 inclusion under hypoxia in TNBC cells. A-C) A) qPCR analysis of alternative splicing of TACC3 B) alternative splicing analysis of TACC3 C) ChIP-qPCR of H3K18la and H3K18ac at CBSDS loci of TACC3 gene region in MCF7 cells with ectopically expressed either empty vector or LDHB in normoxic and hypoxic condition D-F) D) qPCR analysis of alternative splicing of TACC3 E) alternative splicing analysis of TACC3 F) ChIP-qPCR of H3K18la and H3K18ac at CBSDS loci of TACC3 gene region in MCF7 treated with Tamoxifen under hypoxia G-H) G) alternative splicing analysis of TACC3 H) ChIP-qPCR of H3K18la and H3K18ac at CBSDS loci of TACC3 gene region in ESR1 knockdown and shControl in MCF7 cells under normoxia and hypoxia I-J) I) isoform level quantification of TACC3-ENST00000313288 (FL) and TACC3-ENST00000612220 (Δ3 isoform) using SALMON inferred from RNA-seq performed in MDA-MB-231 cells treated with either vehicle (veh) or LDHB inhibitor (LDHBi) under hypoxia J) alternative splicing analysis of TACC3 K) ChIP-qPCR of H3K18la and H3K18ac at CBSDS loci of TACC3 gene region in MDA-MB-231 cells treated with either vehicle (veh) or LDHB inhibitor (LDHBi) under hypoxia. Data are mean ± SEM, statistical significance was determined using one-way ANOVA with Tukey’s multiple comparisons test (A, F, H), unpaired two-tailed Student’s t-tests (B, E, J), ns non-significant, *p < 0.05, **p < 0.01, and ***p < 0.001; n = 3 biological replicates.

LDHB overexpression also shifted H3K18 from the lactylated state in control cells to the acetylated state under hypoxia (**Figure 11C**). We performed the same set of experiments in Tam-treated and ERα-depleted MCF7 cells, both of which increase LDHB expression in MCF7 cells. In agreement with the preceding results, Tam treatment and ERα knockdown each promoted FL isoform production and a switch from H3K18la to H3K18ac in MCF7 cells under hypoxia (**Figure 11D-H**).

We next analysed TACC3 splicing in MDA-MB-231 cells in the absence of LDHB activity. Treatment of MDA-MB-231 cells with LDHBi reduced exon 3 inclusion relative to control cells under hypoxia inferred from RNA-seq and semi-qPCR analysis of TACC3 splicing in MDA-MB-231 cells treated with LDHB inhibitor under hypoxia (**Figure 11I and J**) and shifted H3K18 from the acetylated state in control cells to the lactylated state under hypoxia (**Figure 11K**). Taken together, these findings indicate that LDHB-dependent control of H3K18 lactylation/acetylation status, acting together with CTCF-mediated Pol II pausing, directs the divergent splicing outcomes of TNBC and non-TNBC cells under hypoxia.

### CTCF-mediated kinetic coupling and H3K18la/ac-mediated splicing adaptor recruitment together direct the TACC3 splicing outcome under hypoxia

The dynamics of H3K18la and H3K18ac in TNBC and non-TNBC cells implicate these marks in the regulation of TACC3 splicing under hypoxia. It remained unclear, however, how CTCF-mediated Pol II pausing and H3K18 marks act together to determine the outcome. To address the underlying mechanism of the CTCF-chromatin-splicing adaptor system, we systematically evaluated the determinants of TACC3 splicing.

### CTCF-H3K18la-SMARCA4-hnRNPA1 complex drives TACC3 exon 3 exclusion under hypoxia in non-TNBC cells

As described above, CTCF and H3K18la are each required for the other’s occupancy at CBSDS, and both are required for TACC3 exon 3 exclusion under hypoxia. To understand how they act together, we hypothesized that TACC3 splicing is determined by coupling of CTCF-mediated Pol II pausing (the kinetic model) ^12,27^ with H3K18la-dependent recruitment of splicing factors to chromatin (the adaptor model) ^13^. We searched for chromatin-binding proteins that recognize H3K18la and identified SMARCA4 as a specific reader of this mark ^29^. ChIP-qPCR showed increased SMARCA4 occupancy at CBSDS under hypoxia, which was reduced upon CTCF depletion under hypoxia (**Figure 12A and Figure S5A**), and SMARCA4 depletion increased the FL isoform under hypoxia (**Figure 12B**).

**Figure 12:**
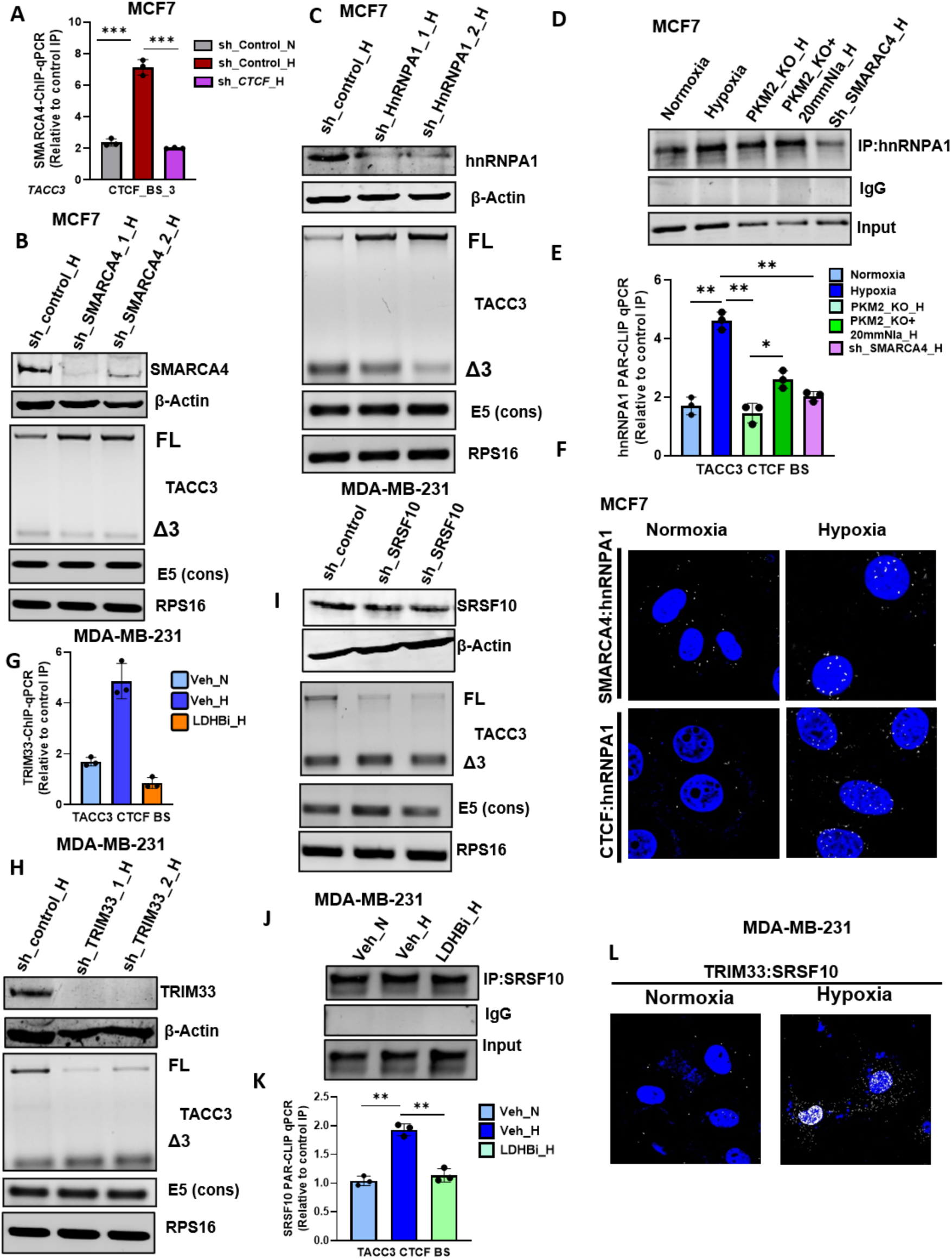
CTCF-mediated kinetic coupling and H3K18la/ac-mediated splicing adaptor recruitment together direct the TACC3 splicing outcome under hypoxia. A) SMARCA4-ChIP-qPCR at TACC3 CBSDS in CTCF knockdown and shControl in MCF7 cells under hypoxia B) immunoblot of SMARCA4 and alternative splicing analysis of TACC3 in SMARCA4 knockdown and shControl in MCF7 cells under hypoxia C) immunoblot of hnRNPA1 and alternative splicing analysis of TACC3 in hnRNPA1 knockdown and shControl in MCF7 cells under hypoxia D) immunoblot of hnRNPA1 in hnRNPA1 pulldown upon PAR-CLIP E) hnRNPA1 PAR-CLIP-qPCR on TACC3 RNA at CBSDS loci in SMARCA4 knockdown and shControl in MCF7 cells under hypoxia F) Representative images of SMARCA4-hnRNPA1 and CTCF-hnRNPA1 PLA (white) 10 hr post in MCF7 under hypoxia G) TRIM33-ChIP-qPCR at TACC3 CBSDS in MDA-MB-231 cells treated with either vehicle (veh) or LDHB inhibitor (LDHBi) under hypoxia H) immunoblot of TRIM33 and alternative splicing analysis of TACC3 in TRIM33 knockdown and shControl in MDA-MB-231 cells under hypoxia I) immunoblot of SRSF10 and alternative splicing analysis of TACC3 in SRSF10 knockdown and shControl in MDA-MB-231 cells under hypoxia J) immunoblot of SRSF10 in SRSF10 pulldown upon PAR-CLIP K) SRSF10 PAR-CLIP-qPCR on TACC3 RNA at CBSDS loci in MDA-MB-231 cells treated with either vehicle (veh) or LDHB inhibitor (LDHBi) under hypoxia L) Representative images of TRIM33-SRSF10 PLA (white) 10 hr post in MDA-MB-231 under hypoxia. Data are mean ± SEM, statistical significance was determined using one-way ANOVA with Tukey’s multiple comparisons test (A, E, G, K), ns non-significant, *p < 0.05, **p < 0.01, and ***p < 0.001; n = 3 biological replicates.

This indicates that hypoxia-induced, CTCF-associated H3K18la recruits SMARCA4 to CBSDS and promotes TACC3 exon 3 exclusion. Because chromatin reader proteins bind specific histone modifications and can in turn interact with and recruit splicing factors to their target genes ^13^, we next asked which splicing factor SMARCA4 recruits to drive exon 3 exclusion. We identified an hnRNPA1 binding site near CBSDS and examined its involvement in TACC3 splicing. hnRNPA1 depletion increased exon 3 inclusion in TACC3 mRNA under hypoxia (**Figure 12C**), and hnRNPA1 binding to TACC3 transcripts at CBSDS was reduced upon SMARCA4 depletion under hypoxia (**Figure 12D-E and Figure S5C-D**). Proximity ligation assay (PLA) confirmed interaction of SMARCA4 with hnRNPA1 and of CTCF with hnRNPA1 under hypoxia (**Figure 12F**). Together, these data indicate that increased CTCF and H3K18la occupancy at CBSDS recruits SMARCA4, which in turn recruits hnRNPA1 to drive TACC3 exon 3 exclusion under hypoxia in non-TNBC cells.

### A CTCF-H3K18ac-TRIM33-SRSF10 complex drives TACC3 exon 3 inclusion under hypoxia in TNBC cells

As described above, CBSDS in TNBC cells is occupied by both CTCF and H3K18ac, and this is associated with inclusion of exon 3 in TACC3 mRNA under hypoxia. We therefore examined the mechanism driving exon 3 inclusion in TNBC cells under hypoxia. Screening for readers of H3K18ac identified TRIM33 as a direct binder of this mark ^30^. TRIM33 occupancy at CBSDS increased under hypoxia and was reduced upon treatment with LDHBi (**Figure 12G and Figure S5E**), and TRIM33 depletion in MDA-MB-231 cells shifted splicing towards exon 3 exclusion under hypoxia (**Figure 12H**). We then asked which splicing factor mediates exon 3 inclusion. SRSF10 depletion shifted splicing towards exon 3 exclusion in MDA-MB-231 cells under hypoxia (**Figure 12I**). SRSF10 binding to TACC3 transcripts at CBSDS increased under hypoxia relative to normoxic control and was reduced upon LDHBi treatment (**Figure 12J-K and Figure S5F-G**). PLA revealed a strong interaction between TRIM33 and SRSF10 under hypoxia that was reduced upon LDHBi treatment (**Figure 12L and Figure S5H**). Together, these data indicate that increased CTCF and H3K18ac occupancy at CBSDS recruits TRIM33, which in turn recruits SRSF10 to drive TACC3 exon 3 inclusion under hypoxia in TNBC cells.

### TACC3 FL isoform drives stemness while Δ3 isoform drives EMT under hypoxia

From the above results, the TACC3 splicing outcome differs between TNBC and non-TNBC cells under hypoxia, with TNBC cells favouring exon 3 inclusion. We next examined the functional roles of the two isoforms, FL and Δ3 TACC3.

We first examined the function of the FL isoform. Overexpression of the FL isoform increased mammosphere formation, whereas the Δ3 isoform had no effect on mammosphere formation, in both MCF7 and MDA-MB-231 cells under hypoxia (**Figure 13A-B**). FL isoform overexpression increased *OCT4* expression in MDA-MB-231 cells and *ALDH1A3* expression in both MCF7 and MDA-MB-231 cells under hypoxia (**Figure 13C-D**), indicating that inclusion of exon 3 under hypoxia promotes stemness. We also observed that the FL isoform resides in the cytoplasm whereas the Δ3 isoform translocates to the nucleus (**Figure 13E**). Overexpression of the FL isoform did not affect the invasiveness of either MCF7 or MDA-MB-231 cells under hypoxia (**Figure 13F**), whereas overexpression of the Δ3 isoform increased invasiveness in both MDA-MB-231 and MCF7 cells under hypoxia (**Figure 13F**) and increased 3D invasion in MCF7 cells (**Figure 13G**). The nuclear localization of the Δ3 isoform may underlie its effect on invasion, potentially through a non-canonical nuclear function that upregulates EMT markers, although this remains to be established. Overall, the FL isoform confers stemness while the Δ3 isoform promotes EMT under hypoxia (**Figure 13H**). This is consistent with the observation that MDA-MB-231 cells already maintain a basal, post-mesenchymal CD44+/CD24-phenotype in which hypoxia can directly activate stemness, whereas MCF7 cells must first acquire a mesenchymal phenotype through EMT ^31^.

**Figure 13:**
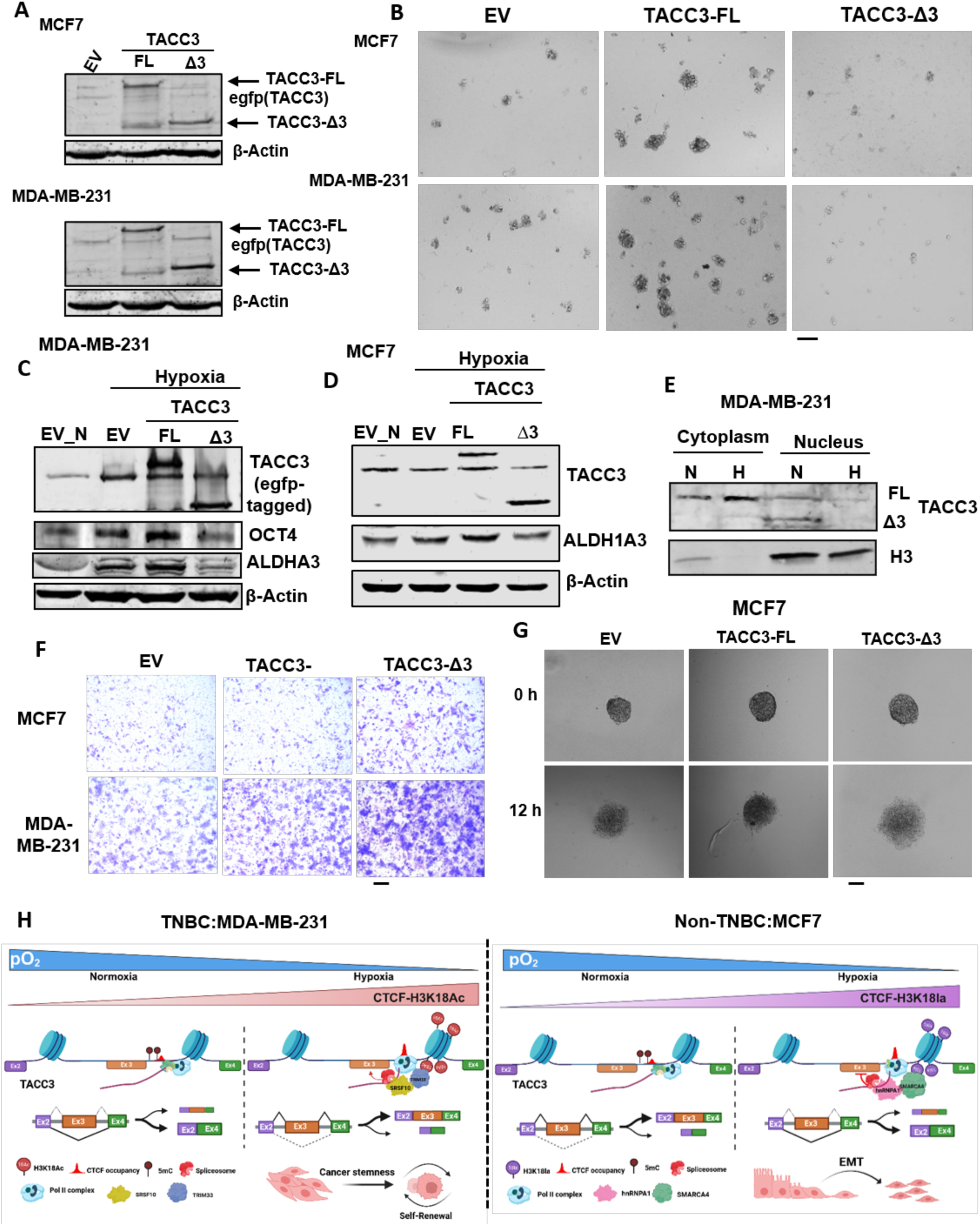
TACC3 FL isoform drives stemness while Δ3 isoform drives EMT under hypoxia. A-B) A) Immunoblot of GFP (to confirm overexpression of TACC3 isoform) B) mammosphere formation assay in MCF7 and MDA-MB-231 cells with ectopically expressed either empty vector or TACC3-FL or TACC3 Δ3 isoform C) Immunoblot of GFP (to confirm overexpression of TACC3 isoform), OCT4 and ALDH1A3 in MDA-MB-231 cells with ectopically expressed either empty vector or TACC3-FL or TACC3 Δ3 isoform D) Immunoblot of GFP (to confirm overexpression of TACC3 isoform), ALDH1A3 in MCF7 cells with ectopically expressed either empty vector or TACC3-FL or TACC3 Δ3 isoform E) immunoblot of TACC3 in cytoplasmic and nuclear fraction of MDA-MB-231 cells under normoxic and hypoxic condition F) invasion assay after overexpression of EV, or TACC3-FL or TACC3 Δ3 isoform in MCF7 and MDA-MB-231 under hypoxia G) Representative images (scale bars, 275 mm) of 3D spheroid invasion assay in MCF7 cells overexpressing EV, TACC3-FL or TACC3 Δ3 isoform spheroids under normoxic conditions H) schematic representation of TACC3 splicing model in Non-TNBC and TNBC and their subsequent effect on cancer phenotype under hypoxia.

### LDHB inhibition reduces tumour growth and stemness in vivo in TNBC

The data presented above indicate that LDHB activity is associated with TACC3 splicing, and that LDHB inhibition in MDA-MB-231 cells decreases inclusion of the TACC3 FL isoform that confers stemness. To test the role of LDHB in stemness, we treated MDA-MB-231 cells with LDHBi under hypoxia and observed reduced expression of *ALDH1A3* and *CD44* (**Figure 14A**), whereas overexpression of LDHB in MCF7 cells increased *ALDH1A3* expression under hypoxia (**Figure 14B**). Hypoxia increased mammosphere formation relative to normoxic MDA-MB-231 cells, and this increase was blunted when LDHBi was given to the hypoxic cells (**Figure 14C**), whereas overexpression of LDHB in MCF7 cells increased mammosphere formation under hypoxia (**Figure 14D**).

**Figure 14:**
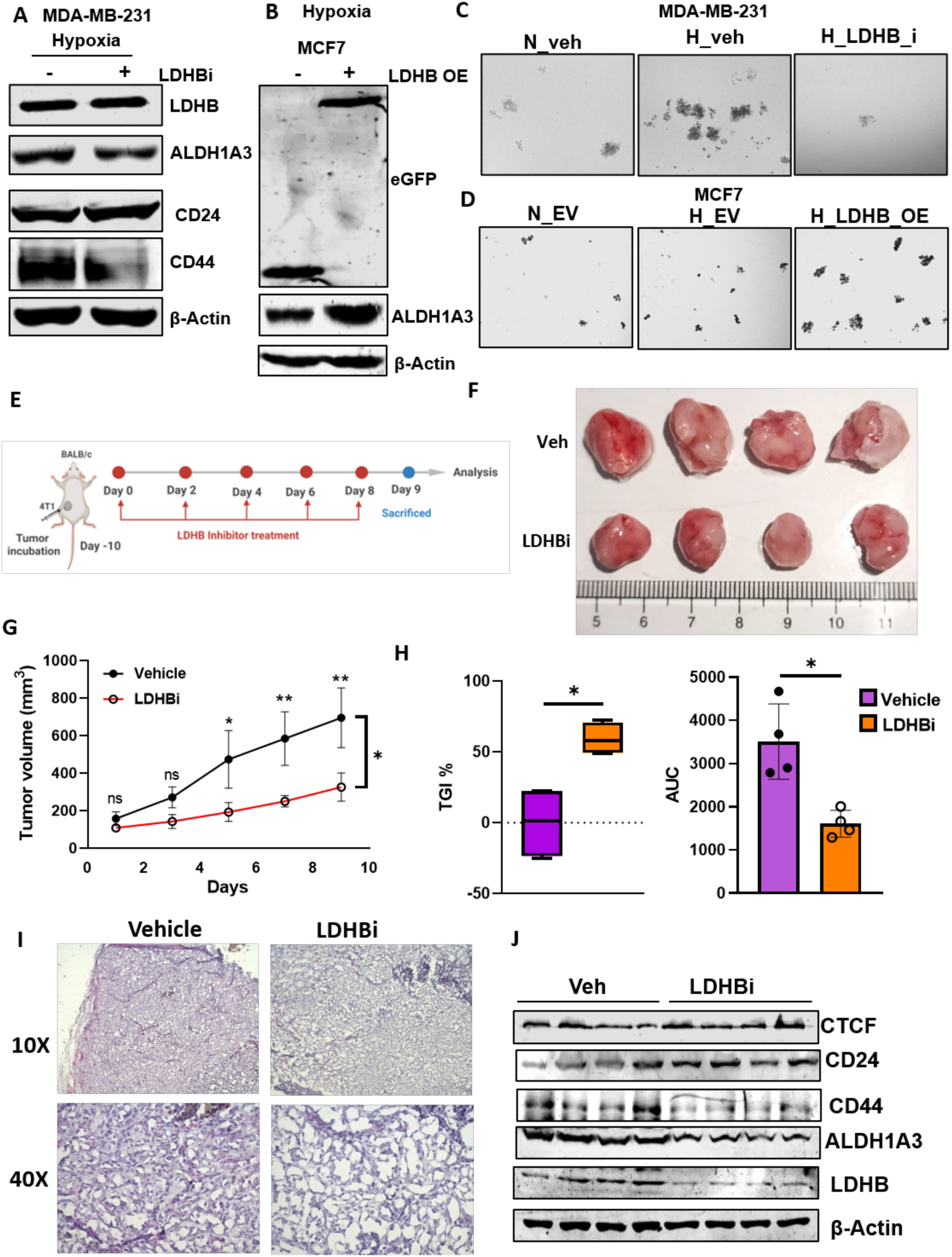
LDHB inhibition reduces tumour growth and stemness in vivo in TNBC. A) Immunoblot of LDHB, ALDH1A3, CD44 and CD24 in MDA-MB-231 cells treated with either vehicle (veh) or LDHB inhibitor (LDHBi) under hypoxia B) Immunoblot of GFP (to confirm overexpression of LDHB) and ALDH1A3 in MCF7 cells with ectopically expressed either empty vector or LDHB in hypoxic condition C-D) mammosphere formation assay C) in MDA-MB-231 cells treated with either vehicle (veh) in normoxia and hypoxia or LDHB inhibitor (LDHBi) under hypoxia D) in MCF7 cells with ectopically expressed either empty vector in normoxia and hypoxia or LDHB in hypoxic condition E) schematic of mouse study. BALB/c mice were inoculated in fat pad with 2 × 10^5^ 4T1 cells in 1:1 RPMI and GFR Matrigel. 11 days after tumor implantation, mice were divided randomly into control and treated group (n=4 mice per group) and either administered with either corn oil or 10 mg/kg of LDHB inhibitor respectively, with oral gavage every alternate day F-I) F) tumor images G) tumor volume H) Tumor growth inhibition I) representative H&E staining J) immunoblot of CTCF, LDHB, CD44, CD24 and ALDH1A3 in control and LDHB inhibitor treated mouse. Data are mean ± SEM, statistical significance was determined using Two-Way Repeated Measures ANOVA Results (G), ns non-significant, *p < 0.05, **p < 0.01, and ***p < 0.001; n = 3 biological replicates unless otherwise indicated.

To test the impact of the LDHB-TACC3 axis on tumour growth, and whether it can be explained by TACC3 splicing, we injected 4T1 cells (a triple-negative murine breast cancer cell line) into BALB/c mice. Mice were assigned to a control group (vehicle) and an LDHBi-treated group (**Figure 14E**). LDHBi significantly reduced tumour volume and tumour growth, supporting a role for LDHB in tumour growth (**Figure 14F-I**). Further analysis revealed reduced expression of *CD44* and *ALDH1A3* and increased expression of *CD24* in LDHBi-treated mice as compared to control mice (**Figure 14J**).

Collectively, our findings indicate that LDHB-dependent epigenetic changes at the TACC3 splicing locus favour exon 3 inclusion and thereby promote stemness, and that LDHB inhibition reverses these stemness phenotypes. These data support the conclusion that disruption of LDHB-mediated lactate metabolism is a promising therapeutic strategy to reduce stemness in TNBC, a subtype that is highly aggressive and prone to chemotherapy resistance.

## Discussion

Metabolic reprogramming and epigenetic rewiring are two central strategies by which cancer cells exploit the adversities imposed by the tumour microenvironment, including nutrient deprivation, acidosis and hypoxia ^32^ ^33^. CCCTC-binding factor (CTCF) is well established as an architectural regulator of three-dimensional genome topology, chromatin looping, enhancer-promoter insulation and transcriptional domain boundaries ^34^, but its direct contribution to the transcription of metabolic genes under physiological and pathological stress has remained largely unexplored. Here we identify CTCF as a transcriptional and epigenetic determinant of metabolic reprogramming that is coupled to alternative splicing, driving divergent outcomes in different breast cancer subtypes under hypoxia (**Figure 15**). CTCF drives glycolytic reprogramming in breast cancer by binding the promoters of key metabolic genes, including *SLC2A3* (GLUT3) and *PFKFB3,* thereby potentiating glucose uptake and glycolytic flux and leading to lactate accumulation under hypoxia. We further define a lineage-dependent regulatory axis in which CTCF controls *LDHB* expression differently in estrogen receptor-positive (ER+) and triple-negative breast cancer (TNBC) models, through functional cooperation with ERα (ESR1).

**Figure 15:**
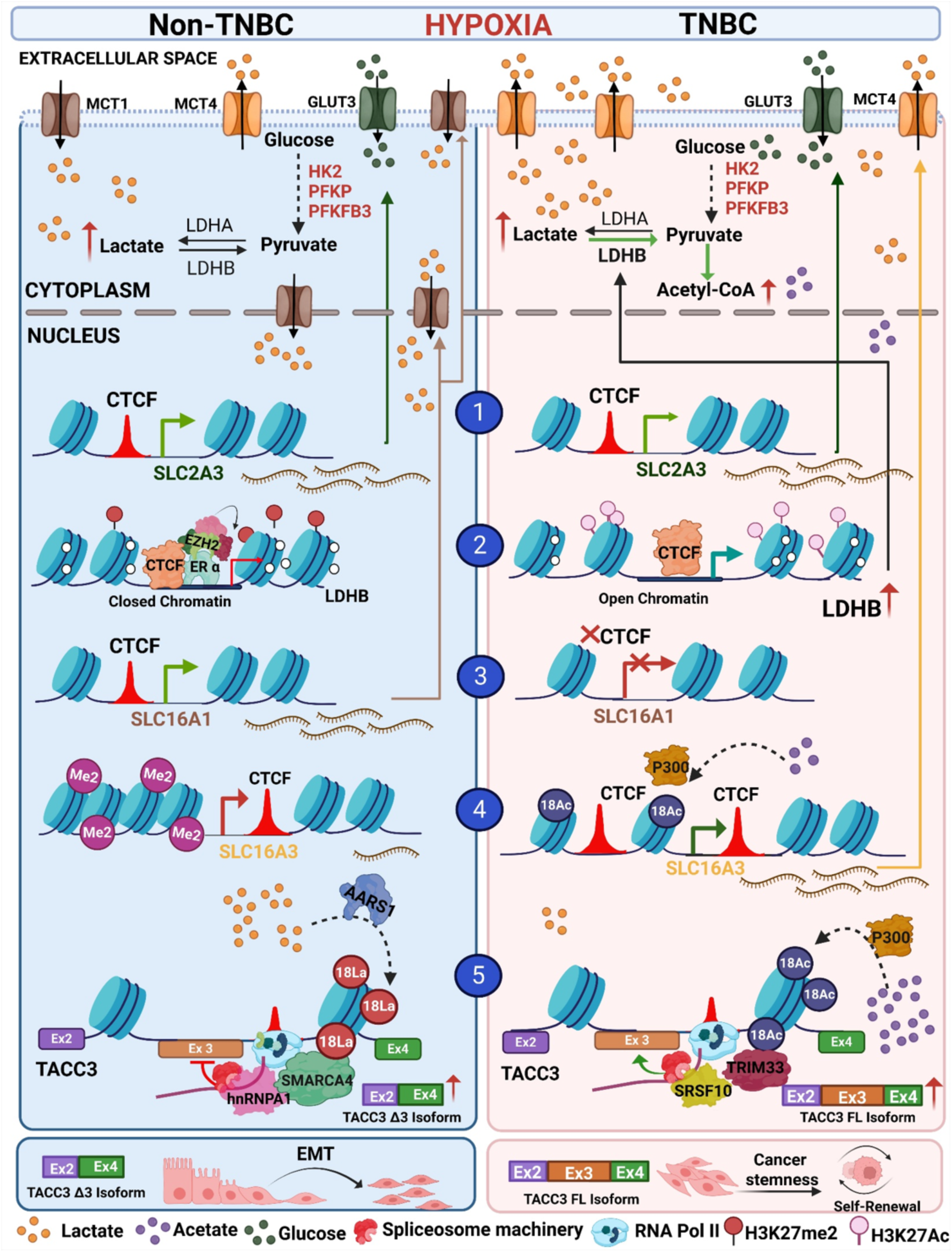
Nuclear lactate availability couples the CTCF-LDHB axis to H3K18 lactylation/acetylation and co-transcriptional splicing in hypoxic breast cancer. Hypoxia bridges metabolic adaptation and chromatin-level transcriptional control in cancer, enhancing the expression of glycolytic genes and thereby supplying both energy and metabolic intermediates for anabolic pathways. Our integrated CTCF ChIP-seq, ATAC-seq and RNA-seq analyses show that hypoxia induces marked chromatin remodelling and increased CTCF occupancy at the promoters of core glycolytic genes. CTCF knockdown under hypoxia significantly reduced SLC2A3 and PFKFB3 expression, accompanied by a decreased extracellular acidification rate (ECAR) and lower intracellular lactate levels. CTCF therefore acts not merely as a passive structural scaffold during hypoxic stress but also promotes local chromatin accessibility, evident from ATAC-seq, and a transcriptionally favourable epigenetic state (H3K27ac) at the promoters of key glycolytic genes, enabling efficient activation of the glycolytic programme to meet the metabolic demands of cancer cells.

A major finding of this study is the contrasting regulation of lactate dehydrogenase B (LDHB) between non-TNBC (MCF7) and TNBC (MDA-MB-231) cell lines. LDHB catalyses the interconversion of lactate and pyruvate, thereby governing lactate oxidation, pyruvate availability and intracellular pH homeostasis, and forming a critical node in the metabolic network. We found that CTCF directly occupies the LDHB promoter in both subtypes, but that the transcriptional consequence is contingent on the activity status of ERα. In MCF7 cells, CTCF and ERα co-occupy the LDHB promoter and expression is repressed, whereas CTCF depletion or tamoxifen (Tam)-mediated inhibition of ERα reduces ERα promoter occupancy and perturbs the repressive histone methylation status (H3K27me2). In contrast, TNBC cells, which lack ERα expression, display a distinct CTCF-dependent epigenetic landscape (H3K27ac) and high LDHB expression. This was further supported by luciferase reporter assays incorporating site-directed mutagenesis of the CTCF and ERα-responsive promoter elements, which attenuated LDHB promoter-driven transcription, particularly in the ER-positive context. This co-regulatory mechanism illustrates how nuclear receptor signalling intersects with architectural factor binding to establish subtype-specific metabolic phenotypes in breast cancer.

Alongside the contrasting expression of LDHB between the two subtypes, we mechanistically define the roles of MCT1 and MCT4 in setting the lactate levels that dictate epigenetic status. MDA-MB-231 cells display a higher extracellular acidification rate (ECAR) than MCF7 cells. In MCF7 cells, CTCF maintains *MCT1* expression and its localization to the nuclear membrane, increasing nuclear lactate and thereby favouring H3K18la at downstream intragenic sites. In MDA-MB-231 cells, increased LDHB expression drives the metabolic shift and, in conjunction with CTCF, favours H3K18ac at the MCT4 promoter, resulting in increased lactate export and reduced nuclear lactate levels.

Our findings demonstrate that, in hypoxic breast cancer cells, metabolic reprogramming redistributes CTCF genome-wide and establishes a subtype-specific metabolic landscape governed by the CTCF-ERα-LDHB axis. A central finding of this work is that the inverse relationship between LDHB and nuclear lactate creates divergent metabolic-epigenetic states across the two breast cancer subtypes, and that these states shape intragenic histone modifications. CTCF, acting as a coordinator of the local chromatin environment, then couples this to co-transcriptional alternative splicing at the downstream target locus TACC3. In MCF7 cells, low LDHB expression together with MCT1-dependent nuclear lactate delivery elevates the local lactate concentration, which favours H3K18 lactylation (H3K18la) at the intragenic CTCF binding site downstream of exon 3. In MDA-MB-231 cells, the absence of ERα permits CTCF-mediated LDHB expression, favouring a metabolic state that drives H3K18 acetylation (H3K18ac) at the identical intragenic CTCF site. CTCF therefore associates with distinct post-translational modifications on adjacent histones in the two subtypes. Both H3K18la and H3K18ac coexist with a CTCF-imposed Pol II pause downstream of TACC3 exon 3, but this kinetic elongation delay is translated into opposite splicing choices depending on which reader protein and splicing factor are recruited.

In non-TNBC cells, H3K18la serves as a binding platform for the SMARCA4 chromatin remodelling complex, which recruits the heterogeneous nuclear ribonucleoprotein hnRNPA1. hnRNPA1 binds a consensus splicing silencer motif near exon 3 and, in the context of the CTCF-imposed Pol II pause, drives exon 3 skipping. The resulting TACC3 isoform lacking exon 3 shifts cellular priorities towards enhanced motility, matrix degradation and invasive capacity under hypoxic stress. Conversely, in TNBC cells the same site is dominated by H3K18ac, which recruits the reader protein TRIM33. TRIM33 serves as an adaptor for the serine/arginine-rich splicing factor SRSF10, which stabilizes 3’/5’ splice site recognition, overrides the default skipping and promotes exon 3 inclusion. Inclusion of exon 3 stabilizes the interactions required for cancer stem cell (CSC) properties, anchorage-independent growth and self-renewal. This mechanism illustrates how breast cancer cells, according to subtype, use intragenic CTCF binding and metabolic-epigenetic crosstalk to toggle TACC3 alternative splicing between invasion and stemness, and defines an adaptor-kinetic coupling model in which CTCF and the local epigenetic landscape together govern co-transcriptional alternative splicing.

## Acknowledgments

P.K. is a recipient of a post-doctoral fellowship from the Indian Institute of Science Education and Research Bhopal, India. B.J., A.B., A.P., and A.S. are recipients of funding from the Indian Institute of Science Education and Research Bhopal, India. Anamika B. is a recipient of a fellowship from the Department of Biotechnology, India. This work is supported by ANRF (ANRF/ARG/2025/007703/LS) and ICMR (IIRPSG-2025-01-01950) (to S.S.).

## Author Contributions

S.S. conceptualized and designed the study. P.K., A.Br. and B.J. designed and performed most of the experiments and analysed the data. R.M.P., A.P., A.Ba., O.P.K., K.N. and A.Sah. performed experiments and contributed to data analysis. P.K. and O.P.K. performed the computational analyses of single-cell, ChIP–seq, ATAC–seq and RNA–seq data. A.Sam. provided clinical input and clinical samples. P.K. and S.S. wrote the manuscript with input from all other authors.

## Data availability

All the sequencing data generated in this study (RNA-seq, all ChIP–seq and ATAC-seq) have been deposited in the Gene Expression Omnibus and will be made publicly available upon publication. This paper does not report original code.

## Material and Methods

### Cell Culture and Hypoxia Treatment

Human breast cancer cell lines MCF-7 and MDA-MB-231, the murine breast cancer cell line 4T1, and the human embryonic kidney cell line HEK293T were obtained from the American Type Culture Collection (ATCC). MCF-7 and HEK293T cells were maintained in Dulbecco’s modified Eagle’s medium (DMEM; Invitrogen, Cat. No. 12800017), 4T1 cells in Roswell Park Memorial Institute (RPMI) medium (Invitrogen, Cat. No. 23400021), and MDA-MB-231 cells in Leibovitz’s L-15 medium (Gibco, Cat. No. 41300039). The culture media were supplemented with 10% FBS (Sigma, Cat. No. F7524) and the reported penicillin–streptomycin and L-glutamine supplements. Cells were maintained at 37°C in a humidified incubator. MCF-7, 4T1, and HEK293T cells were cultured under 5% CO₂, whereas MDA-MB-231 cells maintained in L-15 medium were cultured without supplemental CO₂. Cell line identity was verified by short tandem repeat (STR) profiling through a national cell repository, and cultures were routinely screened for mycoplasma contamination using a PCR-based assay. Hypoxic exposure was performed using a Baker Ruskinn InvivO₂ 400 hypoxia chamber maintained at 1% O_2_ ^9^.

### Animal Experiments

All animal experiments were conducted in accordance with institutional guidelines for the care and use of laboratory animals and were approved by the Institutional Animal Ethics Committee (IAEC). Female BALB/c mice, 4 weeks of age, were used to establish a syngeneic breast tumor model. 2 × 10⁵ murine 4T1 breast cancer cells suspended in 100 µL of a 1:1 mixture of cell suspension and Matrigel were injected orthotopically into the mammary fat pad using a 25-gauge needle. Tumor development was monitored following inoculation, and treatment was initiated on day 11, when tumors had reached approximately 100 mm³. Mice were randomly assigned to vehicle-control and LDHB inhibitor treatment groups. The LDHB inhibitor AXKO-0046 (Cat No. HY-147216) was prepared in a DMSO-containing vehicle supplemented with corn oil and administered by oral gavage at 10 mg/kg body weight in a final volume of 100 µL per dose. Treatment was given every other day for a total of five doses. Tumor dimensions and body weight were recorded on the designated measurement days using a digital vernier caliper and a weighing balance, respectively. Mice were euthanized at the experimental endpoint, before tumors reached the IAEC-approved maximum volume of 1,000 mm³. Tumors were excised, photographed alongside a scale reference, and processed for subsequent molecular and histological analyses. Samples intended for RNA and DNA extraction were snap-frozen, while separate tumor portions were fixed in 4% paraformaldehyde (PFA) for histological examination.

Tumor volume was estimated using the formula (length×width^2^)/2. Tumor growth was assessed by plotting tumor volume against the corresponding experimental day. The area under the tumor growth curve (AUC) was calculated separately for each mouse using the trapezoidal method, and AUC values were compared between the vehicle-control and LDHB inhibitor-treated groups. Data were expressed as mean ± standard error of the mean (SEM), unless otherwise specified. Statistical comparisons were performed using an appropriate two-group test based on the distribution of the data, with *P* < 0.05 considered statistically significant.

### Breast Cancer Patient Samples and Immunoblotting

Breast cancer tissue samples were obtained from Bansal Hospital, Bhopal, India, following written informed consent from the patients. The study was approved by the Institute Ethics Committee of the Indian Institute of Science Education and Research Bhopal, India. Tissue samples representing ER-positive and ER-negative breast cancers were analyzed by Western blotting to assess the expression of ERα, LDHB, and H3K18la, as described in the Fig. Patient clinicopathological details, including age, sex, histological diagnosis, and pathological stage, are provided in the relevant supplementary table (**Table S6**).

### Mammosphere Formation Assay

Mammosphere formation was assessed in MDA-MB-231 and MCF-7 breast cancer cells to examine sphere-forming capacity under hypoxia and following LDHB modulation. MDA-MB-231 cells were evaluated under normoxia, hypoxia, and hypoxia with LDHB inhibitor treatment (2 µM). MCF-7 cells were evaluated under normoxia and hypoxia following empty-vector (EV) transfection, as well as under hypoxia following LDHB overexpression. Adherent cells were harvested by trypsinization, washed twice with phosphate-buffered saline (PBS), and resuspended to obtain a single-cell suspension. Aliquots of 200 µL containing 2,000 MDA-MB-231 cells or 1,000 MCF-7 cells were seeded into Thermo Scientific Nunclon Sphera 96-well flat-bottom ultra-low-attachment plates. The serum-free mammosphere medium was supplemented with insulin (5 µg/mL), hydrocortisone (0.5 µg/mL), bovine serum albumin (BSA; 4 mg/mL), epidermal growth factor (EGF; 20 ng/mL), and basic fibroblast growth factor (bFGF; 20 ng/mL). Cultures were maintained at 37°C under normoxic (21% O₂) or hypoxic (1% O₂) conditions, as appropriate for each group.

Mammospheres were allowed to develop for 2–3 days for MDA-MB-231 cells and 5– 7 days for MCF-7 cells. Culture medium was replenished every 48 h. At the end of the assay, mammospheres were imaged using an inverted phase-contrast microscope and quantified. Mammosphere formation efficiency (MFE) was calculated as the number of mammospheres formed divided by the number of cells seeded, multiplied by 100. MFE (%) = (number of mammospheres formed / number of cells seeded) × 100.

### 2D Transwell Invasion Assay

Cell invasion was evaluated using Transwell inserts with 8.0-µm-pore polycarbonate membranes (Corning, Cat. No. 3422) as previously ^9^. The inserts were coated with 100 µL of Matrigel (Corning, Cat. No. 356230) diluted in serum-free medium at a concentration of 0.25% and incubated at 37°C for 3 h to allow the matrix to set. MCF-7 and MDA-MB-231 cells expressing the indicated TACC3 isoform constructs along with EV were resuspended in serum-free medium, and 3 × 10⁴ cells were seeded into each Matrigel-coated upper chamber. Complete medium supplemented with 10% fetal bovine serum (FBS) was added to the lower chamber to serve as a chemoattractant. The plates were incubated at 37°C under normoxic or hypoxic conditions (1% O₂) for 24–48 h. After incubation, non-invading cells were gently removed from the upper membrane surface with a mild PBS wash. Cells that had traversed the membrane were fixed with 4% paraformaldehyde for 10 min and stained with 0.05% crystal violet prepared in 10% methanol in PBS. The inserts were air-dried, and images were acquired at 10× magnification using an Olympus CKX41 inverted microscope (Tokyo, Japan). Five non-overlapping fields were randomly selected from each insert, and invading cells were quantified using Fiji/ImageJ. Results were expressed as the mean ± standard deviation (SD) of three replicates.

### 3D Spheroid Invasion Assay

The invasive capacity of MCF-7 spheroids expressing the TACC3 isoform constructs and empty vector (EV) was assessed as previously ^12^. For which 3 × 10³ cells were seeded into ultra-low-attachment U-bottom 96-well plates (Thermo Scientific Nunclon Sphera) in culture medium supplemented with hydrocortisone (1 µg/mL; Sigma, Cat. No. H0888) and insulin (5 µg/mL; Sigma, Cat. No. I1882). The plates were incubated at 37°C in a humidified atmosphere containing 5% CO₂ for 24 h to facilitate spheroid formation. Collagen I was prepared at a final concentration of 1 mg/mL, and 100 µL was added to each well of a 96-well culture plate and incubated at 37°C for 1 h to allow polymerization. Individual spheroids were then transferred to the collagen-containing wells and incubated for an additional 1 h at 37°C to allow the matrix to set. Culture medium was subsequently added, and spheroids were maintained under the normoxic or hypoxic conditions for 24h to 48h period. Images were captured at 10× magnification using a Thermo Scientific Invitrogen EVOS FL Auto 2 imaging system. The area of invasion was quantified using ImageJ and expressed relative to the initial spheroid area.

### Immunoblotting

Cells were lysed in a urea-based buffer containing 8 M urea, 2 M thiourea, 2% CHAPS, and 1% dithiothreitol (DTT), supplemented with protease and phosphatase inhibitors. The reported inhibitor supplementation included a 1X protease inhibitor cocktail (PIC; leupeptin 10–100 μM, pepstatin 1 μM, EDTA 1–10 mM, AEBSF < 1 mM), phenylmethylsulfonyl fluoride (PMSF; 10 mM), and 5mM phosphatase inhibitors comprising sodium fluoride (NaF), sodium orthovanadate (Na₃VO₄), and β-glycerophosphate (C₃H₉O₆P). Following lysis, the samples were centrifuged, and the supernatants were collected. Protein concentrations were determined, and equivalent amounts of protein were loaded for electrophoretic separation. Separated proteins were transferred onto activated polyvinylidene difluoride (PVDF) membranes. The membranes were incubated with recommended dilutions of primary antibodies overnight at 4°C, followed by incubation with the corresponding secondary antibodies for 1 h. Protein bands were visualized using a LI-COR Odyssey infrared imaging system, and band intensities were quantified using ImageJ. The antibodies used for immunoblotting are listed in the Table (**Table S7**).

### Pyruvate Assay

Intracellular pyruvate levels were measured using a Pyruvate Colorimetric/Fluorometric Assay Kit (BioVision, Cat. No. K609), following the manufacturer’s instructions. Cells were lysed in the assay buffer supplied with the kit, and the extracts were deproteinized using 10-kDa molecular-weight-cutoff spin columns. The resulting samples were processed with the appropriate assay reagents, and absorbance or fluorescence was measured according to the selected detection method. Pyruvate concentrations were calculated using a standard curve.

### RNA Isolation and cDNA Synthesis

Total RNA was isolated from cultured cells using TRIzol reagent (Invitrogen, Cat. No. 15596026) according to the manufacturer’s instructions. RNA concentration was determined using an Eppendorf BioSpectrometer. For cDNA synthesis, total RNA was reverse-transcribed using the PrimeScript 1st Strand cDNA Synthesis Kit (TaKaRa, Cat. No. 6110A), following the manufacturer’s protocol. The resulting cDNA was used as a template for quantitative and semiquantitative PCR analyses.

### Quantitative Real-Time PCR (RT-qPCR)

Gene and exon expression levels were quantified using KAPA SYBR FAST qPCR Master Mix (Sigma, Cat. No. KK4618) on a qTOWER³ touch real-time PCR system, according to the manufacturer’s instructions. The cycle threshold (Ct) values were used to determine relative expression, with RPS16 serving as the reference gene for normalization. Relative expression was calculated using the (2 (Ct control − Ct target). For exon-level expression analysis, target exon expression was normalized to that of a constitutively expressed reference exon, where applicable. Primer sequences used for qPCR are provided in Supplementary Table (**Table S8**).

### Semiquantitative PCR and Alternative Splicing Analysis

Semiquantitative PCR was performed using cDNA as the amplification template and DreamTaq DNA Polymerase (Thermo Fisher Scientific, Cat. No. EP0702), following the manufacturer’s instructions. Gene-specific and exon-junction primers were used to amplify the target transcripts. The resulting PCR products were separated by electrophoresis on 2% agarose gels with ethidium bromide staining. Primer sequences are listed in Supplementary Table.

Alternative splicing was assessed by examining the relative expression of the relevant splice variants using isoform-specific or exon-junction primers. Where applicable, exon-specific expression was normalized to a constitutive reference exon. Primer sequences used for splice-variant analysis are provided in Supplementary Table (**Table S8**).

### shRNA-Mediated Gene Knockdown

Stable knockdown of CTCF, ESR1 (ERα), TET2, TRIM33, SMARCA4, SRSF10, and HNRNPA1 was established using lentiviral vectors expressing gene-specific short hairpin RNAs (shRNAs). shRNA constructs were obtained from the Sigma Mission Human Genome shRNA Library. Lentiviral particles were generated by co-transfecting HEK293T cells with the respective shRNA-expressing plasmid and the packaging and envelope plasmids (Δ8.9 and VSV-G) using polyethylenimine (PEI). Viral supernatants were collected at 24 and 48 h post-transfection and used to transduce the respective target cells in the presence of polybrene (8 µg/mL; Sigma, Cat. No. H9268). Target cells were seeded at a density of 3 × 10⁵ cells per well in six-well plates and allowed to attach for 24 h before transduction. Following transduction, the medium was replaced, and cells were selected with puromycin (1 µg/mL; Sigma, Cat. No. P9620) for 72 h. The selected cells were subsequently maintained under appropriate culture conditions and used for downstream experiments. Knockdown efficiency was validated by quantitative reverse-transcription PCR (RT-qPCR) and/or immunoblotting, as appropriate for each target. The shRNA target sequences are listed in Supplementary Table (**Table S8**).

### PAR-CLIP qRT-qPCR

Photoactivatable ribonucleoside-enhanced crosslinking and immunoprecipitation (PAR-CLIP) was performed as described previously, with minor modifications. Briefly, cells were supplemented with 100 µM 4-thiouridine (4SU; Cayman Chemical, Cat. No. T4509) 14–16 h before UV crosslinking. Cells were irradiated with 365-nm UV light at a total energy of 0.15 J/cm² using a UV crosslinker, harvested, and lysed in NP-40 lysis buffer (50 mM HEPES-KOH, pH 7.5, 150 mM KCl, 2 mM EDTA-NaOH, pH 8.0, 1mM NaF, and 0.5% NP-40) supplemented with protease inhibitor cocktail, 10 mM PMSF, 0.5 mM DTT, and 40 U/mL RNaseOUT (Invitrogen, Cat. No. 10777019). Lysates were incubated on ice for 10 min and clarified by centrifugation. The supernatants were incubated overnight at 4°C with the appropriate target-specific antibody or control rabbit IgG pre-conjugated to Dynabeads Protein G (Invitrogen, Cat. No. 10004D). Immunoprecipitates were washed sequentially with IP wash buffer containing 300 mM KCl and high-salt wash buffer containing 500 mM KCl, both supplemented with 50 mM HEPES-KOH (pH 7.5), 0.05% NP-40, 0.5 mM fresh DTT, protease inhibitor cocktail, 10 mM PMSF, and 40 U/mL RNaseOUT. An aliquot (20 µL) of each immunoprecipitated complex was retained for immunoblotting. The remaining complexes were treated with Proteinase K buffer (100 mM Tris-HCl, pH 7.5, 12.5 mM EDTA-NaOH, pH 8.0, 150 mM NaCl, 2% SDS, and 1.2 mg/mL Proteinase K) at 55°C for 30 min to reverse crosslinking and digest proteins. RNA was subsequently extracted using TRIzol reagent according to the manufacturer’s instructions, with GlycoBlue coprecipitant (Invitrogen, Cat. No. AM9515) included where applicable. Purified RNA was reverse-transcribed using random hexamers and the PrimeScript 1st Strand cDNA Synthesis Kit (TaKaRa, Cat. No. 6110A). Quantitative PCR was performed using KAPA SYBR FAST qPCR Master Mix (Cat. No. KK4618). Enrichment of the target RNA in the immunoprecipitated fraction was normalized to the corresponding input using 2^(Ct IP − Ct Input). Relative enrichment was then determined by normalizing the target-antibody signal to that obtained with the IgG control.

### Proximity Ligation assay

Proximity ligation assay was performed using Duolink® in Situ Red Starter Kit Mouse/Rabbit (DUO92101-1KT). Human breast cancer cell lines were seeded at the rate of 10^5 cells per glass coverslip in a 12-well plate. Seeded cells were subjected to 12 hrs hypoxic treatment and then fixed with 4% paraformaldehyde for 15 min. The fixed cells were then given a PBS wash and permeabilized with 0.25% Triton X 100. The permeabilized cells were then blocked using Duolink blocking solution (1x) by adding 60 μl in each cover slip and left at room temperature for 1 hour. The primary antibody combinations of two proteins were prepared in a 1:50 ratio of each antibody using Duolink Antibody Diluent. 80 μl of the respective primary antibody combination dilutions were added to the respective cell line coverslips and left at 4°C for overnight incubation. The next day the anti-rabbit (DUO92002) and anti-mouse (DUO92004) secondary antibodies were diluted in a 1:5 ratio with the provided antibody diluent. The primary antibody is removed, and the coverslip is washed with wash buffer A (DUO82049) for 5 min. 60 μl of the above mixture was then added on each coverslip and incubated in a CO₂-free humidified incubator for 1 hour at 37°C. The ligation mixture is prepared by diluting the 5X ligation buffer in high-purity nuclease-free water in a 1:5 ratio. The ligase enzyme is added in the above mixture just before adding on the coverslip in the ratio of 1:40 by discarding the required 1x ligation buffer and adding the corresponding ligase enzyme. The secondary antibody is removed, and the coverslips are washed using wash buffer A for 5 min. The prepared ligation mixture is added and incubated in a CO₂-free humidified incubator for 30 mins at 37°C. The amplification mixture is prepared by diluting the 5X amplification red (light sensitive) in high-purity nuclease-free water in a 1:5 ratio. 1X polymerase enzyme is added to the diluted mixture in the ratio of 1:80 by discarding the required amount of diluted buffer and replacing it with the polymerase enzyme. The ligation mixture is discarded, and the coverslips are washed with wash buffer A for 5 min. 60 μl of amplification mixture is added on each coverslip and incubated in a CO₂-free humidified incubator for 100 mins at 37°C. After amplification the coverslips are washed with wash buffer B two times for 5 min each and then one time using 0.01X wash buffer B. The coverslips are then mounted on glass slides using 30 μl of Duolink In Situ Mounting Medium with DAPI (DUO82040). Imaging was carried out using an Olympus FV3000 confocal laser scanning microscope equipped with a 100 × Plan Apo N oil-immersion objective (numerical aperture 1.42).

### Fluorescence imaging

Approximately 10^5^ cells were seeded onto coverslips in a 12-well plate and allowed to adhere until 70% confluency and then subjected to hypoxic treatment. Post treatment cells were washed three times with ice-cold PBS and then fixed with 4% paraformaldehyde for 15 min. Fixed cells were washed with PBS three times and permeabilized with 0.1% Triton X 100 for 15 min. Cells were then blocked using 1% BSA (in PBST) for 1 hr. After blocking, primary incubation was done overnight at 4°C using MCT 1 antibody. The next day the coverslips with primary antibody were kept at room temperature for 1 hr and then washed using PBST for three times. The cells were then incubated with secondary antibody Rb-555 at a 1:3000 ratio at room temperature for 1 hr. After secondary treatment the cells were washed using PBST for three times and nuclear stained using DAPI at a 1:3000 ratio for 3 min. Nuclear stained cells were then washed with PBST and mounted on a glass slide using Fluoroshield mounting medium (Sigma). Imaging was carried out using an Olympus FV3000 confocal laser scanning microscope equipped with a 60 × Plan Apo N oil-immersion objective (numerical aperture 1.42).

### Seahorse assay

Extracellular acidification rate (ECAR) and oxygen consumption rate (OCR) were measured using the Seahorse XF HS Mini Analyzer. Approximately 10^5^ cells were seeded into six wells of the eight-well of the Seahorse XFp mini cell culture plate, leaving the first and last well empty. Following hypoxic treatment, cells were then washed and incubated with XF assay medium supplemented with 1 mM sodium pyruvate (Sigma, S8636), 2 mM L-glutamine (Invitrogen, 25030081), and 10 mM glucose (Gibco, A24940-01) for 1 hr in a CO_2_-free incubator at 37°C. ECAR was measured following the addition of rotenone/antimycin A (0.5 μM) and 2-deoxy-D-glucose (2-DG; 50 mM), according to the manufacturer’s protocol. OCR measurements were performed with different sequential drug injections of oligomycin (1.5 μM), FCCP (0.5 μM), and a combination of rotenone/antimycin A (0.5 μM each). All Seahorse assay results were normalized to the total protein content per well. The data represent the mean ± standard deviation (SD) from three independent experiments.

### Intracellular lactate assay

Post-treatment media is discarded, and cells are trypsinized and pelleted. Cells are then resuspended in PBS and counted. 10^6^ cells are then centrifuged and lysed using ice-cold assay buffer provided in the lactate assay kit (MAK570-1KT). Lactate quantifications were then carried out using the deproteinized lysates according to the manufacturer’s instructions. The readings were then taken at room temperature with a microplate reader set to 450 nm optical density.

### Extracellular lactate assay

Media is harvested from wells and centrifuged to collect the cell debris and detached cells. The supernatant is then deproteinized using 10 kDa spin columns (Millipore). The deproteinized lysates were then used for the lactate quantification according to the manufacturer’s instructions. The readings were then taken at room temperature with a microplate reader set to 450 nm optical density.

### Nuclear lactate assay

Cells are scraped in ice-cold PBS and pelleted down at 2250 rpm. The pellet is then resuspended in hypotonic solution (10 mM HEPES pH 7.4, 10 mM KCl, 1.5 mM MgCl₂, and 0.5 mM DTT) to isolate nuclear fraction and resuspending thoroughly for 21 times and passing it through a 25-gauge needle. It is then pelleted down at 2250 rpm in 4°C. It is then washed with 500 μl hypotonic solution, and 10 μl is used for cell counting. It is again pelleted down at 2250 rpm at 4°C. The supernatant is removed, and the pellet is resuspended seven times in lactate assay buffer with 0.1% NP40 and sonicated for 8 cycles (Diagenode Bioruptor Pico) with 30 sec on and off intervals for each cycle. The sonicated product is then pelleted down at 12000 rpm for 10 min, and the supernatant is deproteinized using a 10 kDa filter spin column (Millipore). Following the manufacturer’s instructions, the deproteinized lysates were utilized for the lactate quantification. A microplate reader with an optical density of 450 nm was then used to take the readings at room temperature.

### Cloning strategy

Overexpression constructs of LDHB and ESR1 were generated by cloning the insert between Kpn1 and BamH1 of pEFP-C3 (Addgene 6082-1). TACCF FL and TACC3 Δ3 overexpression constructs were cloned into pEFP-C3 using Sal1 and Sma1. For creating the dcas9-HDAC3 and dcas9-AARS1 construct, the DNMT3A CD was digested out of the dCAS9-DNMT3A-puroV2 plasmid (Addgene, 74407) using BamHI and FseI. Following this, MCS taken from pCMV-3tag1A was annealed and cloned in place of DNMT3A CD to create a base parent vector named dcas9-MCS. Subsequently, the HDAC3 and AARS1 full-length constructs were cloned into the dCAS9-MCS using BamHI and Sal1 to generate the dcas9-HDAC3 and dcas9-AARS1 constructs. Primers used for cloning are listed in Table (**Table S8**).

### dcas9-mediated epigenome editing

Guide RNAs (sgRNA) were selected using the GT-Scan web tool and CRISPick and given in **Table S8**. The selected sgRNAs were annealed and cloned at the BbsI restriction site downstream of the U6 promoter in dCas9-DNMT3A-PuroR_v2 (Addgene, 74,407) or the dcas9-HDAC3 and dcas9-AARS1 construct (This paper). Nontargeting sgRNA containing dCas9-DNMT3A vector (Addgene, 71830) was used along with the dCas9-DNMT3A system as a control. The cloned guide RNA was also cloned into dcas9-HDAC3 and dcas9-AARS1 construct and were then verified by sequencing using the U6–F primer. Cells were seeded at 80% confluency, and transfections were carried out at 1:3 ratio of DNA to Turbofect (Thermo Scientific, Cst No. R0531). Cells were then subjected to hypoxic treatment as required post 12 hrs of transfection and harvested for immunoblotting, MeDIP, or ChIP analysis.

### Luciferase Reporter Assay

LDHB promoter fragment ranging from 2076 bp upstream to 176 bp downstream of TSS was amplified using Genomic DNA (gDNA) obtained from MDA-MB-231 cells as template and primes are listed in **Table S8**. Amplified fragment was then cloned between *KpnI* and *HindIII* restriction sites of the pGL3 basic expression vector (Promega, E1751). MCF7 and MDA-MB-231 cells were seeded in 24 well plate and co transfected with LDHB promoter luciferase construct and pRL-TK renilla luciferase plasmid (Promega, E2MDA-MB-231) using PEI (Polyethyleneimine, 23966-100, Polysciences). Post transfection, the cells were subjected to appropriate hypoxia treatment. The cells were then lysed, and luciferase activity was determined using GloMax-Multi Detection System (Promega) by normalizing to the Renilla luciferase activity.

### Site-directed mutagenesis

To generate CTCF, ER∝ and double mutant binding site mutants in LDHB promoter region, SDM primers containing mutations as mentioned in (**Table S8**) were used. The wild-type −2076 to +176 LDHB promoter luciferase construct was used as a template. Following DpnI digestion and transformation, positive colonies were screened and the presence of intended mutations in the CTCF, ER∝ and both site double mutant confirmed by Sanger sequencing.

### Chromatin immunoprecipitation

ChIP assay was performed as previously ^12^. briefly crosslinking about 10 million cells using 1% formaldehyde and quenched by 0.125M glycine. Crosslinked cells were then scrapped in PBS followed by lysis and sonication. Further crosslinked chromatin was sonicated to an approximate chromatin fragment of ∼100–400 bp and chromatin was immunoprecipitated with CTCF, ERα, LDHB, H3K18la, H3K18ac, H3K27ac, H3K27me2, Pol II, EZH2, p300, SMARCA4, TRIM33 antibody or corresponding control (immunoglobin G) antibody overnight at 4°C followed by addition of ProteinG dynabeads (Invitrogen, Cat No. 10004D). The obtained complex and input DNA were reverse cross-linked in TE buffer with 1% SDS and proteinase K (Invitrogen, 25530049, lot no. 2291039 for overnight) at 65°C. Eluted DNA was then purified using PCR purification kit (QIAGEN, 28106). The immunoprecipitated protein–DNA complexes and 5% input were analyzed by qRT-PCR using KAPA SYBR FAST master mix and specific primer sets given in **Table S8**. IP DNA values were normalized to input using the following formula: 2^(Ct_input − Ct_immunoprecipitation (IP)). Resultant values were subsequently normalized to IgG control IP DNA values. The relative values are represented as mean ± SEM of triplicates.

### Genomic DNA isolation and methylated DNA immunoprecipitation (MeDIP-qPCR)

After treatment, treated and control cells were processed to isolate genomic DNA using gDNA isolation kit (Sigma, G1N70) following manufacturer’s instructions. 5μg of gDNA was sonicated to a fragment size of ∼100–400 bp using Bioruptor Sonicator. The sonicated product was then denatured at 95°C for 10 min and then incubated with 5methyl cytosine antibody and rabbit IgG antibody overnight at 4°C. The antibody/DNA complex was then isolated using Dynabeads Protein G (Invitrogen, Cat No. 10004D). Decrosslinking of the immunoprecipitated complexes was performed in TE buffer with 1% SDS and Proteinase K (Invitrogen, 25,530,049) overnight at 65°C. De-crosslinked DNA and input was purified using a PCR purification kit (QIAGEN, 28,106). 5% input and immunoprecipitated samples were analyzed by qRT-PCR using KAPA SYBR FAST master mix and specific primer sets given in **Table S8**. Normalization was performed to input using the formula: [2^(Ct input – Ct immunoprecipitation (IP))]. Resultant values were subsequently normalized to IgG control IP DNA values. The relative values are represented as mean ± SEM of triplicates.

### RNA-seq

For, RNA-seq total RNA was isolated using PureLink RNA Mini Kit (Invitrogen, 12,183,025). mRNA enrichment was performed using NEBNext Poly(A) mRNA Magnetic Isolation Module (E7490S), prior to library preparation. The libraries were constructed using NEBNext Ultra II RNA Library Prep Kit for Illumina (E7770S). These libraries were performed in Novaseq X plus platform.

### ChIP-Sequencing

ChIP-seq experiment was performed as described previously. Briefly, 10 ng of input ChIP-enriched DNA was used and ChIP-seq libraries were performed using NEBNext Ultra II DNA library Prep Kit for Illumina (NEB, Cat. E7645S) according to the manufacturer’s instructions. These libraries were performed in Novaseq X plus platform.

### ATAC-seq

ATAC-seq was performed as described previously ^36^. Briefly, 50,000 MCF7 cells kept under normoxia and hypoxia post 10 hours were harvested and nucleus was isolated. This was further treated Tn5 Transposase (CST catalogue) for 40 min at 37 °C with shaking at 1000 rpm. The samples were purified using MinElute PCR Purification Kit (Qiagen, catalog no. 28004). Library fragments were amplified for 11 cycles using 2× PCR Master Mix (CST) and illumina primers. The libraries were purified using Agencourt AMPure XP beads (Beckman Coulter, catalog no. A63881). The libraries were sequenced as paired-end (150 bp) on Novaseq X plus platform.

### Bioinformatics Analysis

#### Retrieval of gene list from various databases

Gene sets for Hallmark hypoxia, the Buffa hypoxia signature, and glycolysis were retrieved from MSigDB ^37^. The Eukaryotic Promoter Database (EPD) was used for promoter annotation. Gene set overlaps were analyzed using the Vennerable (v3.1.0.9000) package in R.

### Bedtools Analysis

To identify CTCF binding sites associated with glycolysis-related genes, we used BEDTools (v2.31.1) ^38^ to determine the genomic proximity of differential CTCF peaks to transcription start sites (TSSs) annotated using the EPD database. CTCF peaks were classified into three groups: (i) downstream peaks located within 5 kb of glycolytic gene TSSs, (ii) upstream peaks within 5 kb of TSSs, and (iii) peaks overlapping TSS regions. Differential CTCF-binding peaks identified by DiffBind were used to assess potential associations between CTCF occupancy and glycolysis-related genes under normoxic and hypoxic conditions.

### Single-Cell RNA-seq Analysis

Single-cell RNA-seq data from GSE161529 were processed using Seurat (v5.5.1) ^39^. After quality control and filtering, samples were merged, normalized using LogNormalize, and 3,000 highly variable genes were selected for Principal Component Analysis (PCA) and Uniform Manifold Approximation and Projection (UMAP) clustering. Major cell lineages were annotated using marker-gene expression.

Malignant epithelial cells were identified by integrating epithelial annotations with CopyKAT (v1.2.5) ^40^ aneuploidy classifications. Hypoxia and glycolysis scores were calculated using the Buffa hypoxia signature and Hallmark Glycolysis gene set from MSigDB, respectively, using Seurat’s AddModuleScore function. Cells were categorized into high- and low-score groups using median cutoffs.

Cellular glycolytic activity was quantified using the Hallmark Glycolysis gene set (Molecular Signatures Database, MSigDB ^37^) via Seurat’s ‘AddModuleScorè function to generate the ‘Glycolysis_Scorè. Microenvironmental hypoxia status was assessed using the 52-gene Buffa hypoxia metagene signature (‘Buffa_Hypoxia_Score’).

To isolate cells experiencing robust microenvironmental extremes while minimizing intermediate noise, malignant cells were stratified by their continuous Buffa hypoxia scores. Quantiles were established at the 30th and 70th percentiles to define extreme phenotypic subsets: the top 30% (Hypoxic Extreme) and bottom 30% (Normoxic Extreme), with intermediate cells excluded. To eliminate single-cell dropout artifacts and assess coordinate expression at the biological replicate level, single-cell profiles were aggregated into pseudobulk replicates grouped by sample identifier (‘sample’) and extreme hypoxia status. For each biological sample, mean expression of genes CTCF, LDHB, SLC2A3 and SLC16A3), and mean glycolysis scores were calculated. To ensure statistical stability, pseudobulk replicates were filtered to top 20 high-coverage samples per phenotypic group. Linear relationships between pseudobulk mean CTCF expression and glycolytic scores were evaluated using Pearson correlation analysis independently for the extreme hypoxic and normoxic cohorts. Binarized co-occurrence was further assessed using Fisher’s Exact Test. Statistical significance was established at p< 0.05. All computational analyses and visualizations were executed in R.

### RNA-seq data analysis

Raw sequencing reads in FASTQ format were assessed for quality using FastQC (v0.12.1) ^41^. Raw reads were trimmed for adapters and low-quality bases (Phred score < 20) using fastp (v0.23.4) ^42^. Reads were aligned to the human reference genome (GRCh38 / hg38) using STAR (v2.7.11b) ^43^.

Gene-level read counts were generated using featureCounts (v2.0.6) with the Ensembl GRCh38 (release 111) annotation. Downstream differential expression analysis was conducted using DESeq2 (v1.42.0) ^44^. Low-expressed genes (fewer than 10 total counts across all samples) were filtered out before normalization. Gene annotation was performed using BioMart (v2.58.0) ^45^ to retrieve gene symbols and descriptions. Genes with an adjusted p-value (False Discovery Rate, FDR) < 0.05 and an absolute log2 fold change ≥ 1 were classified as differentially expressed genes (DEGs).

### ChIP-seq data analyses

Reads were aligned to human genome hg38 using Bowtie2 ^46^. Alignments were subsequently filtered for high-confidence mappings using Samtools with a quality threshold of MAPQ ≥ 30 (-q 30) and sorted by genomic coordinate. Peak calling was performed using MACS3 (v3.0.1) with matched input controls. Differential peak analysis of ChIP-seq data was performed using DiffBind (v3.12.0) ^47^. Heatmaps were generated using deepTools (v3.5.6) ^48^ to visualize enrichment across the identified peaks.

Using the ShinyGO (v 8.6) ^49^ web server, Gene Ontology analysis was performed, and pyGenomeTracks was used to visualize the signals.

### ATAC-seq data analyses

Reads were aligned to human genome hg38 using Bowtie2. Peaks were identified using MACS3. Differential chromatin accessibility was assessed by performed using DiffBind (v3.12.0). BigWig files were prepared using and deepTools was used for heatmap generation.

### Isoform Expression and Relative Usage Analysis

Transcript-level expression using Salmon, and transcript abundance estimates were imported using tximport to generate expression matrices for downstream differential expression analysis. Transcript annotations were obtained from the reference GTF file.

Isoform-level expression quantification for the target gene *TACC3* was analyzed using Transcript Per Million (TPM) values. To characterize transcriptional dynamics under normoxic and hypoxic conditions, individual transcript abundance values for the predominant isoforms, TACC3-201 and TACC3-212, were extracted alongside whole-gene expression data. Relative isoform usage was calculated as the percentage of individual isoform expression relative to the total accumulated expression of all 18 detected *TACC3* transcripts within each sample. Low-abundance isoforms with maximum usage of less than 1% across all samples were filtered out, and a hierarchical heat map was constructed using average relative usage metrics to organize and visualize the remaining predominant isoforms.

## References

1. Seyfried, T. N. & Huysentruyt, L. C. On the origin of cancer metastasis. Crit. Rev. Oncog. 18, 43–73 (2013).

2. Pavlova, N. N. & Thompson, C. B. The Emerging Hallmarks of Cancer Metabolism. Cell Metabolism vol. 23 Preprint at 10.1016/j.cmet.2015.12.006 (2016).

3. Muz, B., de la Puente, P., Azab, F. & Azab, A. K. The role of hypoxia in cancer progression, angiogenesis, metastasis, and resistance to therapy. Hypoxia 10.2147/hp.s93413 (2015) doi:10.2147/hp.s93413.

4. Lei, P. et al. Role of Glucose Metabolic Reprogramming in Breast Cancer Progression and Drug Resistance. Cancers vol. 15 Preprint at 10.3390/cancers15133390 (2023).

5. Shin, E. & Koo, J. S. Glucose Metabolism and Glucose Transporters in Breast Cancer. Frontiers in Cell and Developmental Biology vol. 9 Preprint at 10.3389/fcell.2021.728759 (2021).

6. Xiao, S. et al. Lactate and lactylation: molecular insights into histone and non-histone lactylation in tumor progression, tumor immune microenvironment, and therapeutic strategies. Biomarker Research vol. 13 Preprint at 10.1186/s40364-025-00849-0 (2025).

7. Zhang, D. et al. Metabolic regulation of gene expression by histone lactylation. Nature 574, 575–580 (2019).

8. Kindrick, J. D. & Mole, D. R. Hypoxic regulation of gene transcription and chromatin: Cause and effect. International Journal of Molecular Sciences vol. 21 Preprint at 10.3390/ijms21218320 (2020).

9. Kakani, P. et al. Hypoxia-induced CTCF promotes EMT in breast cancer. Cell Rep. 43, 114367 (2024).

10. Yadav, P. et al. Hypoxia-induced loss of SRSF2-dependent DNA methylation promotes CTCF-mediated alternative splicing of VEGFA in breast cancer. iScience 10.1016/j.isci.2023.106804 (2023) doi:10.1016/j.isci.2023.106804.

11. Dhamdhere, S. G. et al. Hypoxia-induced ATF3 escalates breast cancer invasion by increasing collagen deposition via P4HA1. Cell Death Dis. 16, 142 (2025).

12. Kakani, P. et al. Hypoxia-induced CTCF mediates alternative splicing via coupling chromatin looping and RNA Pol II pause to promote EMT in breast cancer. Cell Rep. 44, 115267 (2025).

13. Yustis, J. C., Devoucoux, M. & Côté, J. The Functional Relationship Between RNA Splicing and the Chromatin Landscape. Journal of Molecular Biology vol. 436 Preprint at 10.1016/j.jmb.2024.168614 (2024).

14. Farhana, A. & Lappin, S. L. Biochemistry, Lactate Dehydrogenase (LDH). StatPearls (2020).

15. Mishra, D. & Banerjee, D. Lactate dehydrogenases as metabolic links between tumor and stroma in the tumor microenvironment. Cancers vol. 11 Preprint at 10.3390/cancers11060750 (2019).

16. Jordan, V. C. Tamoxifen: A most unlikely pioneering medicine. Nature Reviews Drug Discovery vol. 2 Preprint at 10.1038/nrd1031 (2003).

17. Margueron, R. et al. Ezh1 and Ezh2 Maintain Repressive Chromatin through Different Mechanisms. Mol. Cell 32, (2008).

18. Hu, Y. et al. Lactylation: the novel histone modification influence on gene expression, protein function, and disease. Clin. Epigenetics 16, 72 (2024).

19. Wang, T., et al. Lactate-induced protein lactylation: A bridge between epigenetics and metabolic reprogramming in cancer. Cell Proliferation vol. 56 Preprint at 10.1111/cpr.13478 (2023).

20. Lv, X., Lv, Y. & Dai, X. Lactate, histone lactylation and cancer hallmarks. Expert Reviews in Molecular Medicine Preprint at 10.1017/erm.2022.42 (2023).

21. Li, X. et al. Lactate metabolism in human health and disease. Signal Transduction and Targeted Therapy vol. 7 Preprint at 10.1038/s41392-022-01151-3 (2022).

22. Navale, A. M. & Paranjape, A. N. Glucose transporters: physiological and pathological roles. Biophysical Reviews vol. 8 Preprint at 10.1007/s12551-015-0186-2 (2016).

23. TeSlaa, T. et al. The Source of Glycolytic Intermediates in Mammalian Tissues. Cell Metab. 33, (2021).

24. Bosshart, P. D., Kalbermatter, D., Bonetti, S. & Fotiadis, D. Mechanistic basis of L-lactate transport in the SLC16 solute carrier family. Nature Communications 10, (2019).

25. Hong, J. et al. AARS1-mediated lactylation of H3K18 and STAT1 promotes ferroptosis in diabetic nephropathy. Cell Death Differ. 33, (2026).

26. Zong, Z. et al. Alanyl-tRNA synthetase, AARS1, is a lactate sensor and lactyltransferase that lactylates p53 and contributes to tumorigenesis. Cell 187, (2024).

27. Shukla, S. et al. CTCF-promoted RNA polymerase II. Nature 10.1038/nature10442 (2011) doi:10.1038/nature10442.

28. Moreno-Yruela, C. et al. Class I histone deacetylases (HDAC1-3) are histone lysine delactylases. Sci. Adv. 8, (2022).

29. Hu, X. et al. Dux activates metabolism-lactylation-MET network during early iPSC reprogramming with Brg1 as the histone lactylation reader. Nucleic Acids Res. 52, (2024).

30. Sun, H. et al. Recruitment of TRIM33 to cell-context specific PML nuclear bodies regulates nodal signaling in mESCs. EMBO J. 42, (2023).

31. Srivastava, N., Usmani, S. S., Subbarayan, R., Saini, R. & Pandey, P. K. Hypoxia: syndicating triple negative breast cancer against various therapeutic regimens. Frontiers in Oncology vol. 13 Preprint at 10.3389/fonc.2023.1199105 (2023).

32. Tsai, Y. P. & Wu, K. J. Epigenetic regulation of hypoxia-responsive gene expression: Focusing on chromatin and DNA modifications. International Journal of Cancer Preprint at 10.1002/ijc.28190 (2014).

33. Camuzi, D. et al. Regulation Is in the Air: The Relationship between Hypoxia and Epigenetics in Cancer. Cells 10.3390/cells8040300 (2019) doi:10.3390/cells8040300.

34. Phillips, J. E. & Corces, V. G. CTCF: Master Weaver of the Genome. Cell Preprint at 10.1016/j.cell.2009.06.001 (2009).

35. Corces, M. R. et al. An improved ATAC-seq protocol reduces background and enables interrogation of frozen tissues. Nat. Methods 14, (2017).

36. Liberzon, A. et al. Molecular signatures database (MSigDB) 3.0. Bioinformatics 27, (2011).

37. Quinlan, A. R. & Hall, I. M. BEDTools: A flexible suite of utilities for comparing genomic features. Bioinformatics 10.1093/bioinformatics/btq033 (2010) doi:10.1093/bioinformatics/btq033.

38. Hao, Y. et al. Dictionary learning for integrative, multimodal and scalable single-cell analysis. Nat. Biotechnol. 42, 293–304 (2024).

39. Gao, R. et al. Delineating copy number and clonal substructure in human tumors from single-cell transcriptomes. Nat. Biotechnol. 39, (2021).

40. Andrews, S. FastQC - A quality control tool for high throughput sequence data. http://www.bioinformatics.babraham.ac.uk/projects/fastqc/. Babraham Bioinformatics (2010).

41. Chen, S. fastp 1.0: An ultra-fast all-round tool for FASTQ data quality control and preprocessing. iMeta 4, (2025).

42. Dobin, A. et al. STAR: Ultrafast universal RNA-seq aligner. Bioinformatics 10.1093/bioinformatics/bts635 (2013) doi:10.1093/bioinformatics/bts635.

43. Love, M. I., Huber, W. & Anders, S. Moderated estimation of fold change and dispersion for RNA-seq data with DESeq2. Genome Biol. 10.1186/s13059-014-0550-8 (2014) doi:10.1186/s13059-014-0550-8.

44. Smedley, D. et al. BioMart - Biological queries made easy. BMC Genomics 10.1186/1471-2164-10-22 (2009) doi:10.1186/1471-2164-10-22.

45. Langmead, B., Trapnell, C., Pop, M. & Salzberg, S. L. Ultrafast and memory-efficient alignment of short DNA sequences to the human genome. Genome Biol. 10.1186/gb-2009-10-3-r25 (2009) doi:10.1186/gb-2009-10-3-r25.

46. Rory, S. & Gord, B. DiffBind: Differential binding analysis of ChIPSeq peak data. Bioconductor (2011).

47. Ramírez, F. et al. deepTools2: a next generation web server for deep-sequencing data analysis. Nucleic Acids Res. 44, W160–W165 (2016).

48. Ge, S. X., Jung, D., Jung, D. & Yao, R. ShinyGO: A graphical gene-set enrichment tool for animals and plants. Bioinformatics 10.1093/bioinformatics/btz931 (2020) doi:10.1093/bioinformatics/btz931.

