## Supplementary Figures for "A CTCF-LDHB axis converts nuclear lactate availability into an H3K18 acylation switch that directs alternative splicing in hypoxic breast cancer"

**Supplementary File**

### Supplementary Figure 1

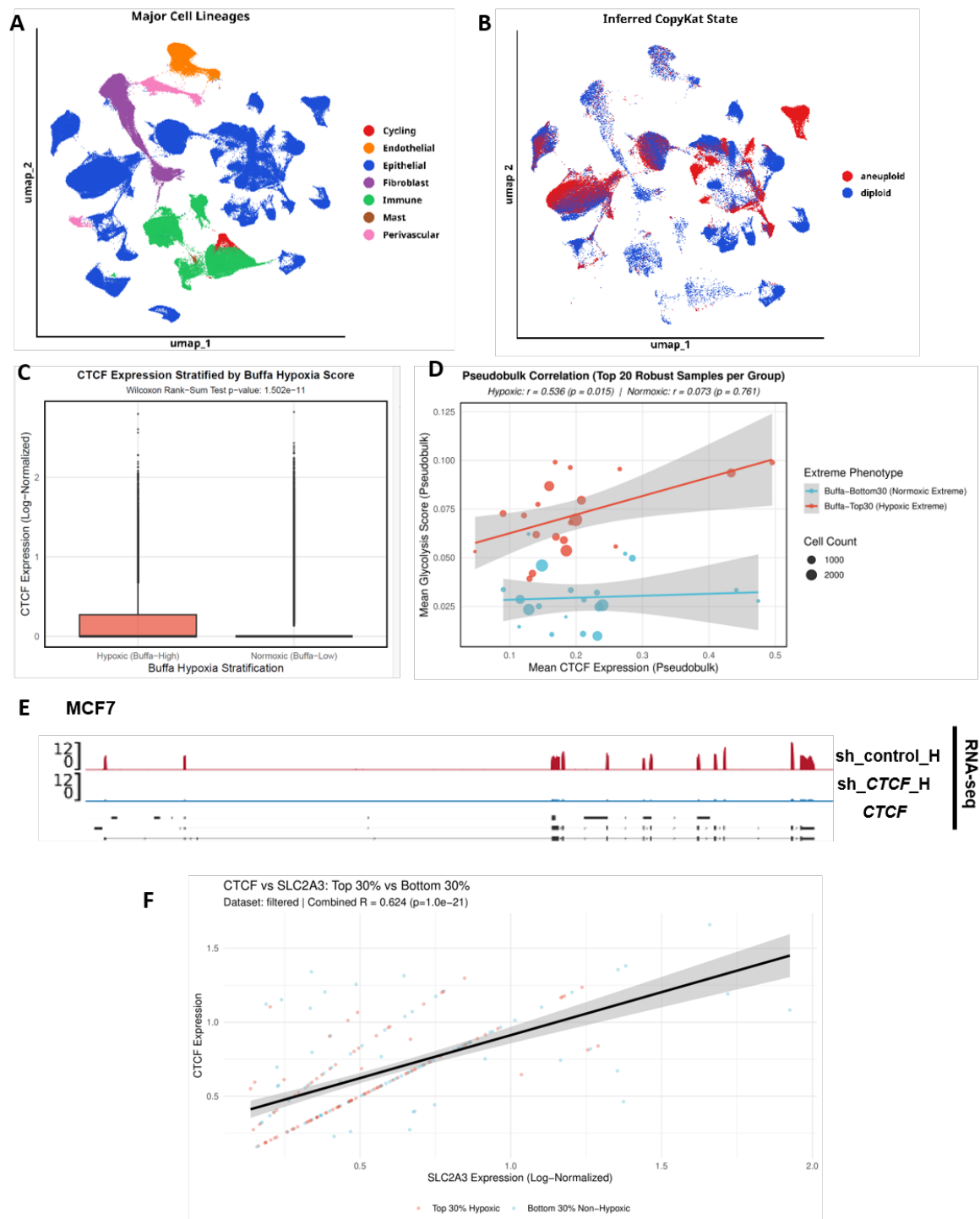

**Supplementary Figure 1:** A) uniform manifold approximation and projection (UMAP) clustering of breast epithelial cancerous cell populations with major cell lineages annotated using established canonical marker genes B) UMAP of malignant epithelial cells identified by integrating lineage annotations with CopyKAT aneuploidy classifications C) CTCF expression levels stratified by extreme Buffa hypoxia score

quantiles in breast cancer single-cell dataset GSE161529. Statistical significance was determined using a two-sided Wilcoxon rank-sum test D) Pseudobulk correlation analysis between mean CTCF expression and mean glycolysis score across extreme Buffa hypoxia groups (top 20 robust samples per group) from dataset GSE161529. Red points and regression line represent the Hypoxic extreme group, while blue points and regression line represent the Normoxic Extreme group E) genome browser view of CTCF loci of RNA-seq in shcontrol and shCTCF under hypoxia in MCF7 cells F) Correlation scatter plot between SLC2A3 and CTCF expression comparing Top 30% Hypoxic and Bottom 30% Non-Hypoxic conditions in the filtered dataset. Red points denote samples in the hypoxic group, while blue points represent samples in the non-hypoxic group.

### Supplementary Figure 2

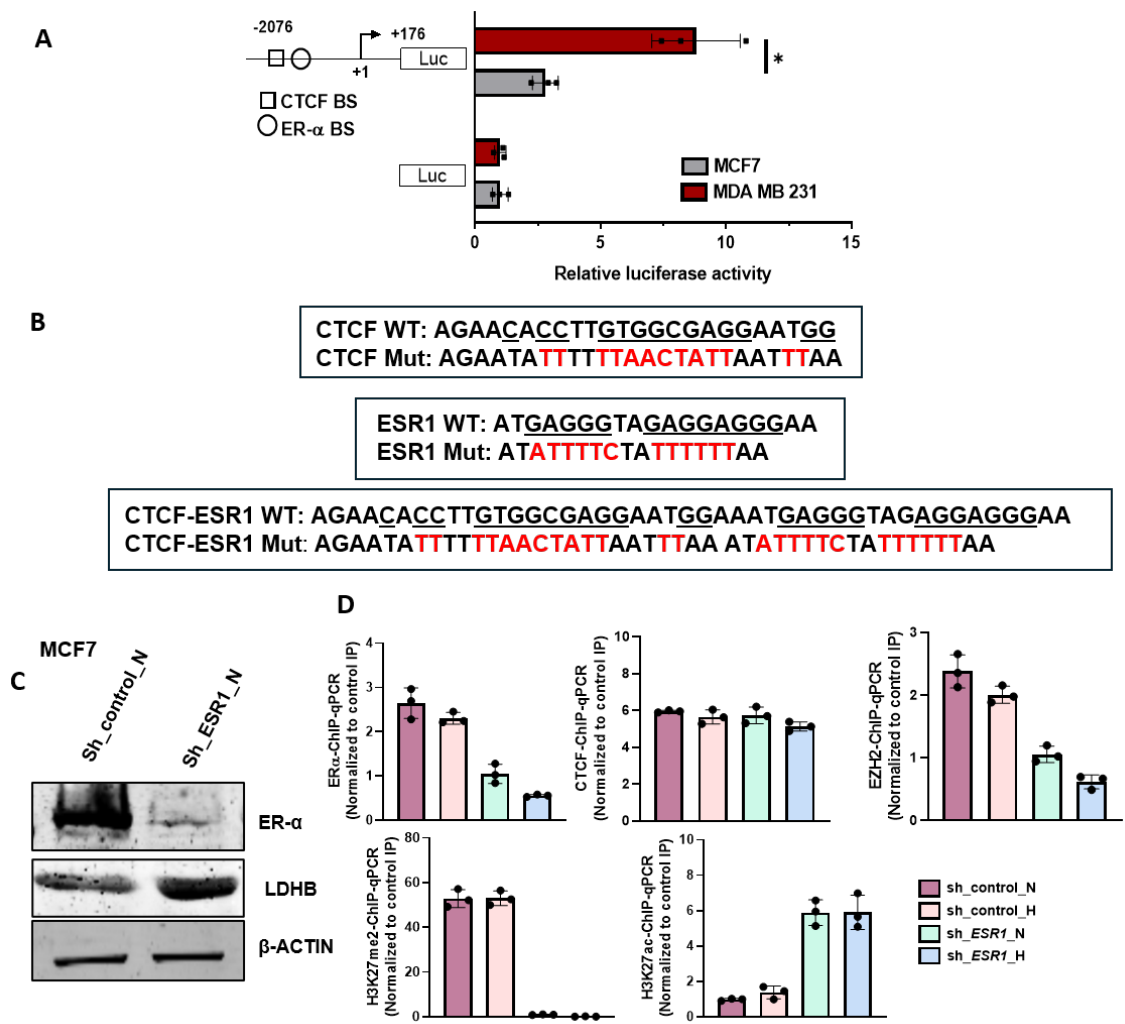

**Supplementary Figure 2:** A) MCF7 and MDA-MB-231 cells were co-transfected with wild type LDHB promoter luciferase construct (-337 to +115) and pRL-TK Renilla under normoxia B) Schematic representation of the wild-type and mutated type or CTCF or ERα or CTCF/ ERα mutant sequences for LDHB promoter deletion constructs C) Immunoblot of ERα and LDHB in ESR1 depletion in comparison to sh\_control in MCF7 cells D) ERα, CTCF, EZH2, H3K27me2 and H3K27ac-ChIP-qPCR on LDHB promoter in ESR1 depletion in comparison to sh\_control in MCF7 cells under normoxia and hypoxia. Data are mean ± SEM, statistical significance was determined using one-way ANOVA with Tukey's multiple comparisons test. ns non-significant, \*$p < 0.05$, \*\*$p < 0.01$, and \*\*\*$p < 0.001$; $n = 3$ biological replicates.

#### Supplementary Figure 3

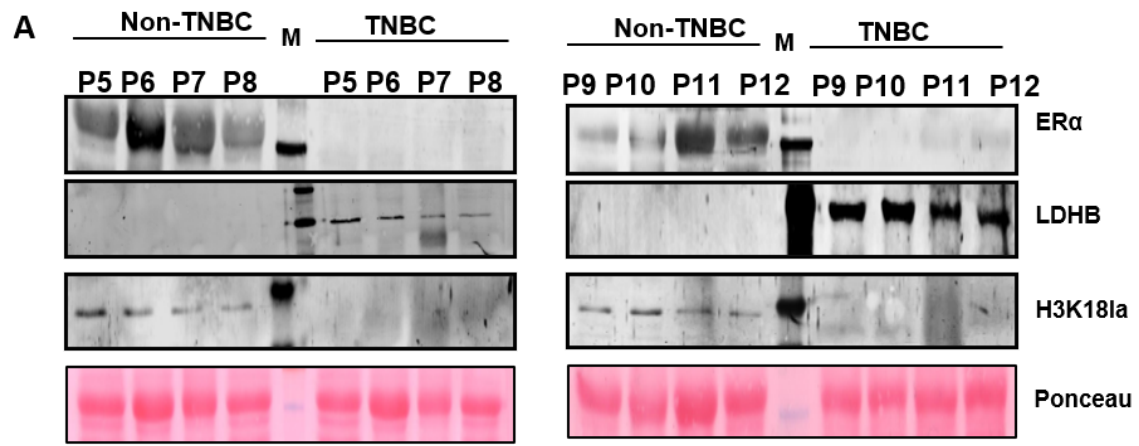

**Supplementary Figure 3:** A) Immunoblot of ERα, LDHB and H3K18la in breast tumor samples of non-TNBC and TNBC patients

### Supplementary Figure 4

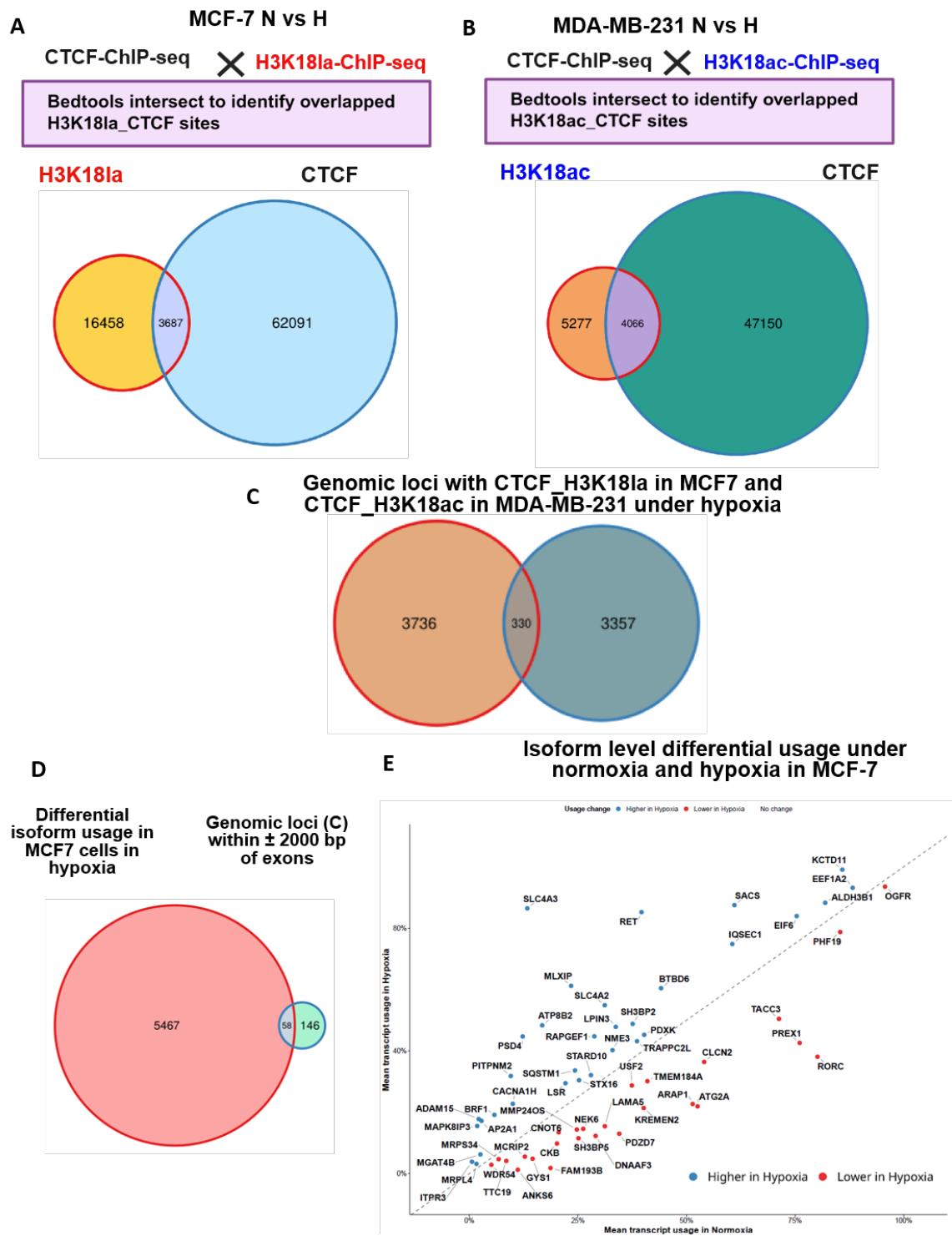

**Supplementary Figure 4:** A) Analysis of H3K18la and CTCF binding sites inferred from ChIP-seq, Venn diagram representing number of genomic sites co-occupied by hypoxia gained H3K18la site and CTCF in MCF7 cells B) Analysis of H3K18ac and CTCF binding sites inferred from ChIP-seq, Venn diagram representing number of

genomic sites co-occupied by hypoxia gained H3K18ac site and CTCF in MDA-MB-231 cells C) Venn diagrams showing the genomic loci occupied by H3K18la-CTCF in MCF7 while H3K18ac-CTCF in MDA-MB-231 cells D) Venn diagram showing overlap between differential isoform usage in MCF7 in hypoxia in comparison to normoxic control inferred from RNA-seq using SALMON and DRIMSeq and genomic loci obtained from (C) that were falling in window of $\pm 2000$ bp of exons (intersected using bedtools with human exon coordinates to locate the H3K18la-CTCF and H3K18ac-CTCF) E) volcano plot differential isoform usage of exons obtained from (D) in MCF7 under normoxia and hypoxia.

### Supplementary Figure 5

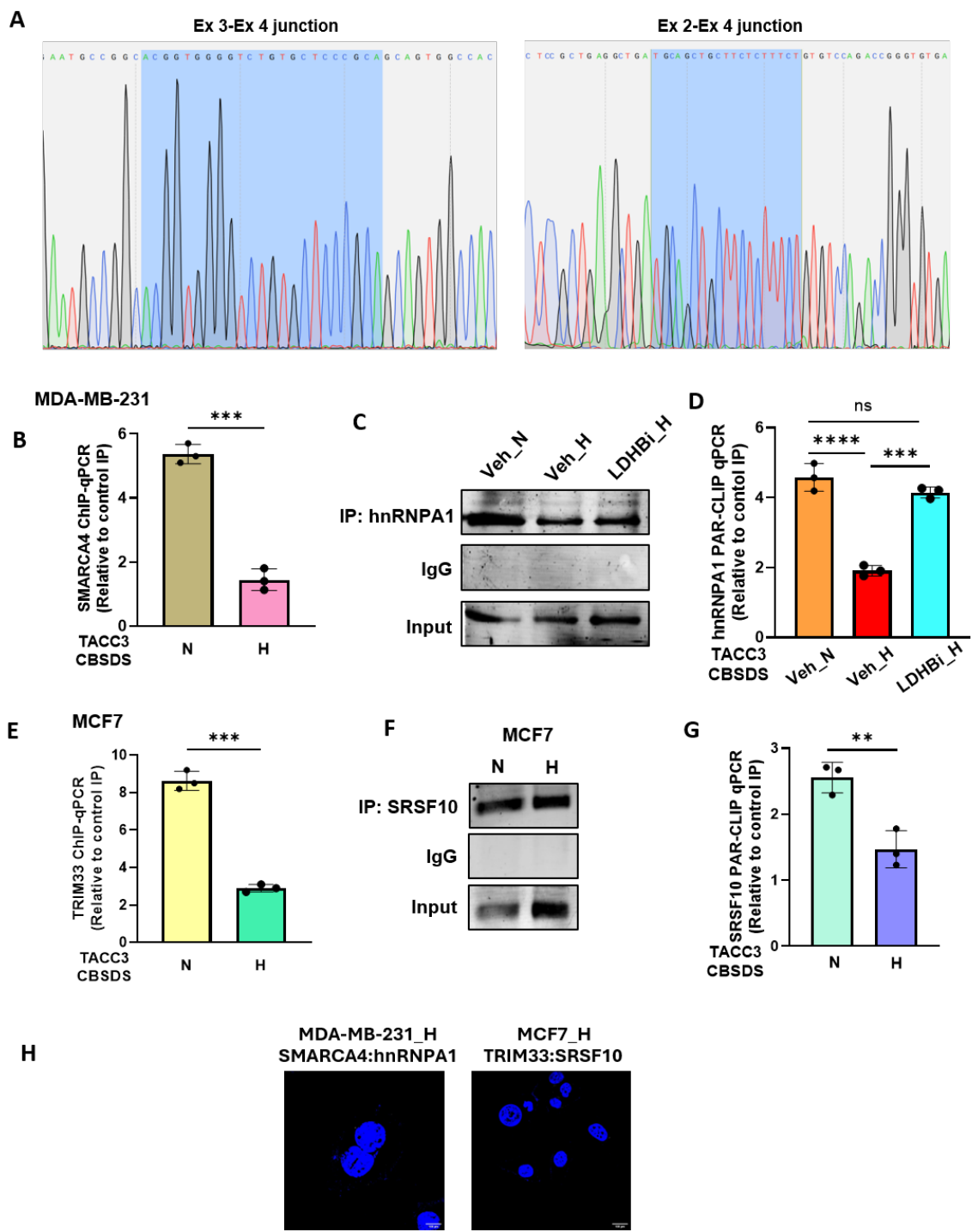

**Supplementary Figure 5:** A) chromatogram representing the TACC3 gene alternative splicing, exon3-4 junction, give rise to TACC3-full length (FL) and exon 2-4 junction, give rise to $\Delta 3$ isoform, B) SMARAC4-ChIP-qPCR at TACC3 CDSDS in MDA-MB-231 cells under normoxia and hypoxia C) immunoblot of hnRNPA1 in hnRNPA1 pulldown upon PAR-CLIP D) hnRNPA1 PAR-CLIP-qPCR on TACC3 RNA at CDSBS loci in

SMARAC4 knockdown and shControl in MDA-MB-231 cells treated with either vehicle (veh) or LDHB inhibitor (LDHBi) under hypoxia E) TRIM33-ChIP-qPCR at TACC3 CDSDS in MCF7 cells normoxia and hypoxia F) immunoblot of SRSF10 in SRSF10 pulldown upon PAR-CLIP G) SRSF10 PAR-CLIP-qPCR on TACC3 RNA at CDSBS loci in MCF7 cells normoxia and hypoxia H) Representative images of SMARCA4-hnRNPA1 and TRIM33-SRSF10 PLA (white) 10 hr post in MDA-MB-231 and MCF7 respectively under hypoxia. Data are mean ± SEM, statistical significance was determined using one-way ANOVA with Tukey's multiple comparisons test (A, E, G, K), ns non-significant, \*$p < 0.05$, \*\*$p < 0.01$, and \*\*\*$p < 0.001$; n = 3 biological replicates.
